# Tripartite chromatin localization of budding yeast Shugoshin involves higher-ordered architecture of mitotic chromosomes

**DOI:** 10.1101/274241

**Authors:** Xiexiong Deng, Min-Hao Kuo

**Affiliations:** Department of Biochemistry and Molecular Biology, Michigan State University. East Lansing, MI 48824

## Abstract

The spindle assembly checkpoint (SAC) is key to faithful segregation of chromosomes. One requirement that satisfies SAC is appropriate tension between sister chromatids at the metaphase-anaphase juncture. Proper tension generated by poleward pulling of mitotic spindles signals biorientation of the underlying chromosome. In the budding yeast, the tension status is monitored by the conserved Shugoshin protein, Sgo1p, and the tension sensing motif (TSM) of histone H3. ChIP-seq reveals a unique TSM-dependent, tripartite domain of Sgo1p in each mitotic chromosome. This domain consists of one centromeric and two flanking peaks 3 – 4 kb away, and is present exclusively in mitosis. Strikingly, this trident motif coincides with cohesin localization, but only at the centromere and the two immediate adjacent loci, despite that cohesin is enriched at numerous regions throughout mitotic chromosomes. The TSM-Sgo1p-cohesin triad is at the center stage of higher-ordered chromatin architecture for error-free segregation.

## INTRODUCTION

Equal partition of the duplicated chromosomes is crucial for genome integrity and species perpetuation. Aneuploidy resulting from erroneous segregation causes developmental defects and tumorigenesis (Ricke et al., 2008). The spindle assembly checkpoint (SAC) is a failsafe for faithful segregation. The SAC registers the kinetochore-microtubule attachment and the tension between sister chromatids (Pinsky and Biggins, 2005). The tension generated by poleward pulling of the spindles signals bipolar attachment, after which cells irreversibly initiates events leading to the onset of anaphase.

In *Saccharomyces cerevisiae*, each kinetochore attaches to a single microtubule spindle emanating from the spindle pole bodies (Cleveland et al., 2003). To the two sister kinetochores, three types of attachment may occur: monotelic, syntelic and amphitelic (Pinsky and Biggins, 2005). While the amphitelic attachment signals biorientation, monotelic and syntelic attachment errors have to be corrected before anaphase onset. Monotelic attachment refers to the situation when only one of the two sister kinetochores is attached to the microtubule. The presence of an unoccupied kinetochore triggers the formation of the Mitotic Checkpoint Complex (MCC) (Brady and Hardwick, 2000) that halts cell cycle progression by trapping Cdc20p, the E3 ligase subunit of Anaphase Promoting Complex (APC). In syntelic attachment, both sister kinetochores are occupied by spindles, but these two spindles originate from the same spindle pole body. Even though the attachment requirement is met, there may be no tension between syntelic sister chromatids as they are pulled toward the same pole. Left uncorrected, monotelic and syntelic attachment results in aneuploidy.

In what form tension is perceived by the mitotic machinery remains elusive. In prometaphase, transient sister chromatid separation without cohesin proteolysis is caused by kinetochore-microtubule attachment (He et al., 2001, He et al., 2000). Conformational changes of centromeric chromatin (DNA, nucleosomal arrays, and selective proteins) thus are suggested to be the “tensiometer” or “spring” that reflects the tension status (Salmon and Bloom, 2017). Among these candidates, Shugoshin proteins are of particular interest. Shugoshin is a family of conserved proteins playing critical roles in ensuring appropriate chromatid cohesion during cell division (Marston, 2015). The budding yeast Shugoshin, Sgo1p, was first identified as a protector of meiotic cohesin against precocious cleavage (Kitajima et al., 2004), and later found to be also crucial for cells to activate the SAC in coping with tensionless conditions in mitosis (Indjeian et al., 2005). Expressed in S and M phases of the cell cycle (Indjeian et al., 2005, Eshleman and Morgan, 2014), Sgo1p is localized to centromeres and pericentromere (Fernius and Hardwick, 2007, Kiburz et al., 2005, Kiburz et al., 2008) without stashing a significant extrachromosomal pool (Buehl et al., 2018). Shugoshin is recruited to centromeres by binding to histone H2A phosphorylated by the Bub1 kinase (Kawashima et al., 2010, Liu et al., 2013a). The centromeric recruitment of budding yeast Sgo1p may also involve the interaction with the centromere-specific histone H3 variant Cse4p (Mishra et al., 2017). In human mitotic cells, Sgo1 recruited to the outer kinetochore nucleosomes is then driven by RNA polymerase II to the inner centromere where it is retained by cohesin (Liu et al., 2015). Besides cohesin, the fission yeast meiosis-specific Shugoshin Sgo1 interacts with the heterochromatin protein 1 (HP1) homologue Swi6 that docks on the heterochromatic mark H3K9me3 in pericentromere (Yamagishi et al., 2008, Isaac et al., 2017). Unlike other eukaryotes where heterochromatic marks decorate pericentromere to create a footing for Shugoshin, budding yeast lacks such heterochromatic features in the region immediately next to centromeres (Cleveland et al., 2003). The geographic pericentromere recruitment of Sgo1p in budding yeast, instead, is accomplished by the association with the tension sensing motif (TSM) of histone H3 in pericentric regions (Luo et al., 2010, Luo et al., 2016). TSM (^42^KPGT) is a conserved β-turn that connects the flexible N’ tail to the rigid histone-fold domain of H3 (White et al., 2001). Mutations at K43, G44, or T45 diminish the pericentric localization of Sgo1p and obliterate the cellular response to defects in tension. Restoring pericentric association of Sgo1p by overexpression, via Sgo1p-bromodomain fusion (Luo et al., 2010), or by mutating the inhibitory residues K14 or K23 of the H3 tail (Buehl et al., 2018) rescues the mitotic defects of these TSM mutations, thus manifesting the pivotal role of Sgo1p retention at the pericentromere. Sgo1p is removed from chromatin after tension is built up in the metaphase (Nerusheva et al., 2014). The inverse correlation between Sgo1p retention and amphitelic attachment suggests that Sgo1p is an integral part of the gauge by which cells use to monitor the tension status.

In addition to the TSM, another factor important for targeting Sgo1p to the pericentromere is the cohesin complex. Mutations that impair cohesin loading ablate pericentric localization of Sgo1p, while leaving the centromeric Sgo1p largely unaffected (Kiburz et al., 2005). A similar contribution of cohesin to Sgo1 localization has been observed in human systems as well (Liu et al., 2015). Cohesin performs it tension sensing-related function by facilitating the formation of the “C” loop of chromatin near the centromeres in mitosis (Stephens et al., 2011, Yeh et al., 2008). Direct interaction between cohesin and the human Sgo1 has been reported (Liu et al., 2013b). The triad of Sgo1, H3 TSM, and cohesin thus likely constitute the core of the tension sensing device. The present work presents evidence for a cohesin- and TSM-dependent tripartite chromatin localization domain of Sgo1p that also involves high-ordered chromatin architecture.

## RESULTS

### Sgo1p displays unique tripartite localization in each mitotic chromosome

Sgo1p is critical for the tension sensing branch of the SAC function in mitosis (Marston, 2015). We and others have previously used chromatin immunoprecipitation (ChIP) to demonstrate that Sgo1p is enriched at centromeres and several kb on either side of the centromere in mitosis (Luo et al., 2010, Kiburz et al., 2005, Nerusheva et al., 2014, Fernius and Hardwick, 2007). To better understand Sgo1p retention pertaining to its checkpoint function, we used ChIP-seq to map the Sgo1p distribution on mitotic chromosomes at a higher resolution. Cells bearing a C-terminally HA-tagged Sgo1p expressed from its native locus were arrested by benomyl for ChIP-seq. At a lower resolution scale, Sgo1p is detectable in one area per mitotic chromosome (Supplemental Figure 1A), consistent with the anticipation of centromeric and pericentric enrichment (Kiburz et al., 2005). However, more rigorous inspection revealed that each chromosomal domain of Sgo1p is actually composed of discrete peaks of Sgo1p that form a trident-like structure, not a continuous motif covering several kb of a centromeric and pericentric area (Figure 1). Each of the trident motif consists of a middle centromere (CEN) and typically one pericentromere (PC) peak on each side of the CEN enrichment. By aligning all sixteen chromosomes at the centromeres, the average counts plot for Sgo1p enrichment as a function of distance to CEN shows that the average distance between the PC and CEN peaks is approximately 4 kb (Figure 2A, magenta line). Additional outward peaks may be seen in some chromosomes but the overall peak height drops quickly.

**Figure 1.**
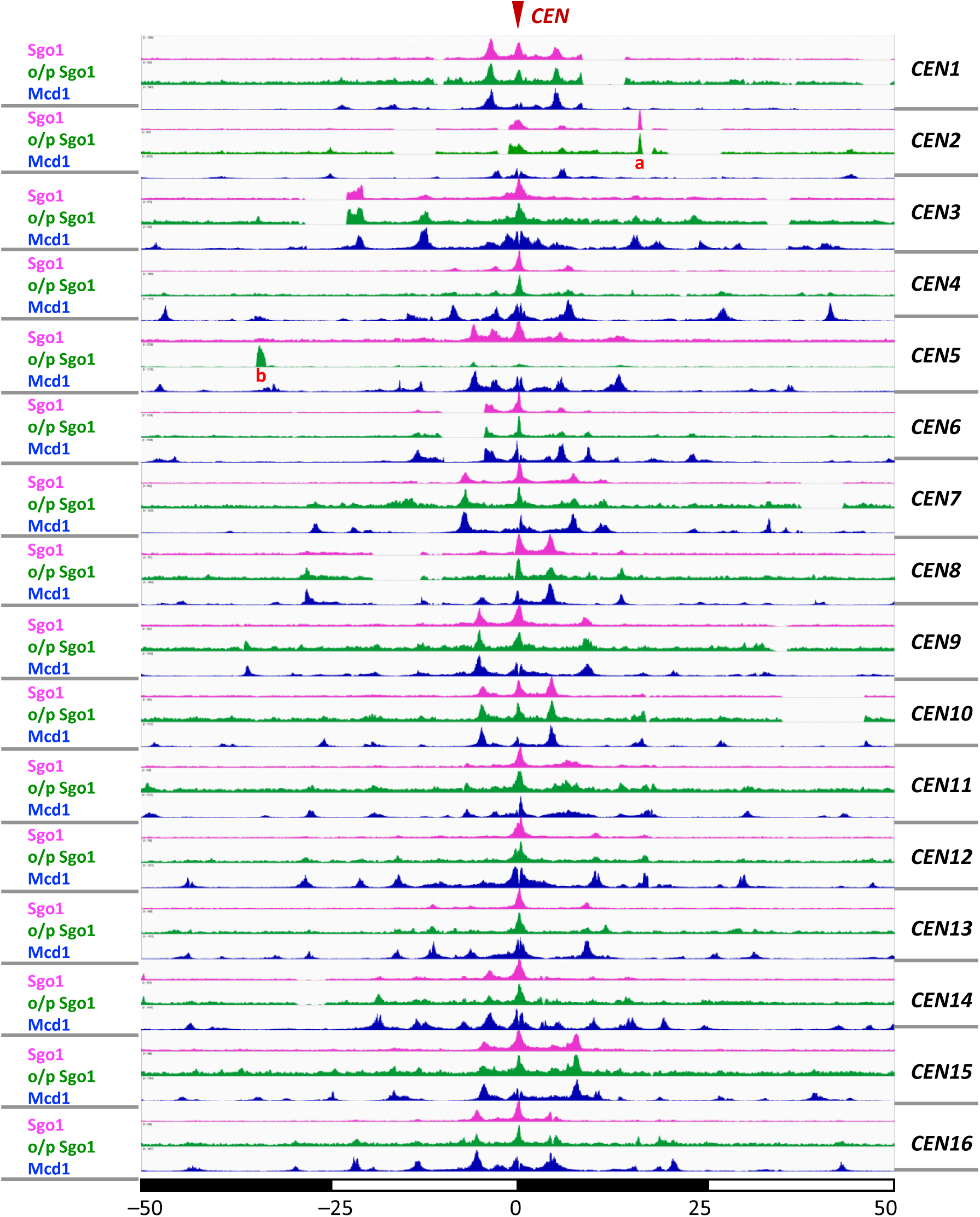
Sgo1p is recruited to centromeres and pericentromere to form a tripartite localization domain on each mitotic chromosome. The 100-kb region centering on the centromere of all 16 chromosome is aligned. Sgo1p expressed from its native locus (magenta), or from a multi-copy episomal plasmid (green) are compared with the Mcd1p distribution (dataset from Verzijlbergen et al., 2014). The two peaks labelled “a” and “b” close to *CEN2* and *CEN5* correspond to *ARS209* and *URA3* respectively. These loci were from two plasmids in the strains used for experiments.

**Figure 2.**
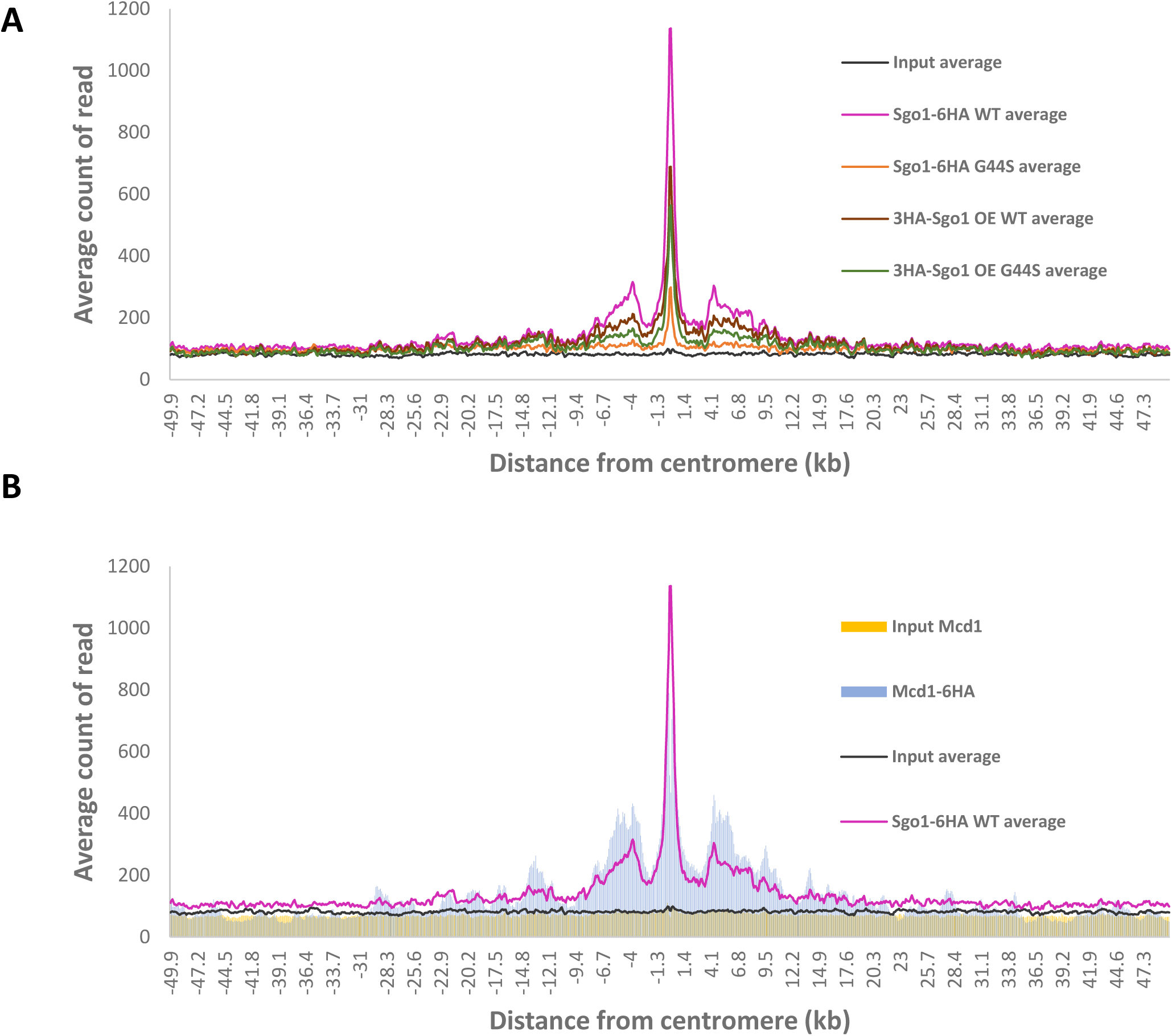
Sgo1p enrichment overlaps with cohesin domains at the centromeres and pericentromere. Average counts (per million reads) plot comparing the distribution of Sgo1p expressed in different backgrounds (panel A) or between Sgo1p and cohesin (panel B). The counts apparently corresponding to plasmid-borne *URA3* and *ARS209* were artificially removed from the calculation. The Mcd1p ChIP-seq data were from Verzijlbergen et al., 2014.

Chromosomal retention of Sgo1p depends critically on the tension sensing motif (TSM) of histone H3 (Luo et al., 2010), and the cohesin complex (Verzijlbergen et al., 2014). H3 is a ubiquitous component of chromatin, yet it controls the pericentric localization of Sgo1p (Luo et al., 2010), despite that no discernible epigenetic marks have been found specifically in budding yeast pericentromere that are relevant to mitotic regulation. Mutations introduced to the tension sensing motif (^42^KGPT^45^) cause defects in detecting and/or responding to tension defects (Luo et al., 2016). These mutations diminish the affinity for Sgo1p, a molecular defect that can be suppressed by overproduction of Sgo1p (Luo et al., 2016, Luo et al., 2010). Indeed, ChIP-seq data show that the overall chromatin association of Sgo1p is significantly reduced in a tension sensing motif mutant, G44S (Supplemental Figure 1A, orange curve). Expressing Sgo1p from a multi-copy plasmid and the *ADH1* promoter restored the tripartite chromatin association (green curves, Figure 1, and brown curve, Supplemental Figure 1A). In addition to re-establishing the original enrichment pattern, a small number of new peaks distal to the CEN/PC peaks were seen. Intriguingly, these still are discrete peaks with clear valleys in between (see, for example, chromosome XVI, Figure 1). The emergence of these new enrichment is consistent with our original model that Sgo1p is recruited to the centromeres and then spills over to the nearby chromatin region (Luo et al., 2010). However, the non-continuous nature of Sgo1p distribution suggests the involvement of at least one other factor (see below).

While histone H3 and its tension sensing motif are ubiquitously distributed throughout the genome, another Sgo1p recruitment factor, the cohesin complex, localizes at specific loci of chromosomes. Besides centromeres and pericentric regions, the majority of cohesin-associated regions are the intergenic region between two convergent transcription units throughout the genome (Lengronne et al., 2004, Glynn et al., 2004). By comparing with the chromosomal distribution of Mcd1p (the kleisin subunit of cohesin) (Verzijlbergen et al., 2014), we observed that Sgo1p co-localizes with cohesin at and immediately adjacent to centromeres (compare magenta and blue peaks, Figure 1 and Figure 3A). The plot of average count reads (Figure 2B) clearly shows the highly significant co-localization of Sgo1p- and Mcd1p at the centromeric and pericentric region. It is also noteworthy that most additional Sgo1p peaks resulting from overexpression are at the loci where cohesin is also enriched (Figure 1). These results strongly suggest that Sgo1p targets existing cohesin enrichment sites for interaction with the tension sensing motif of histone H3.

**Figure 3.**
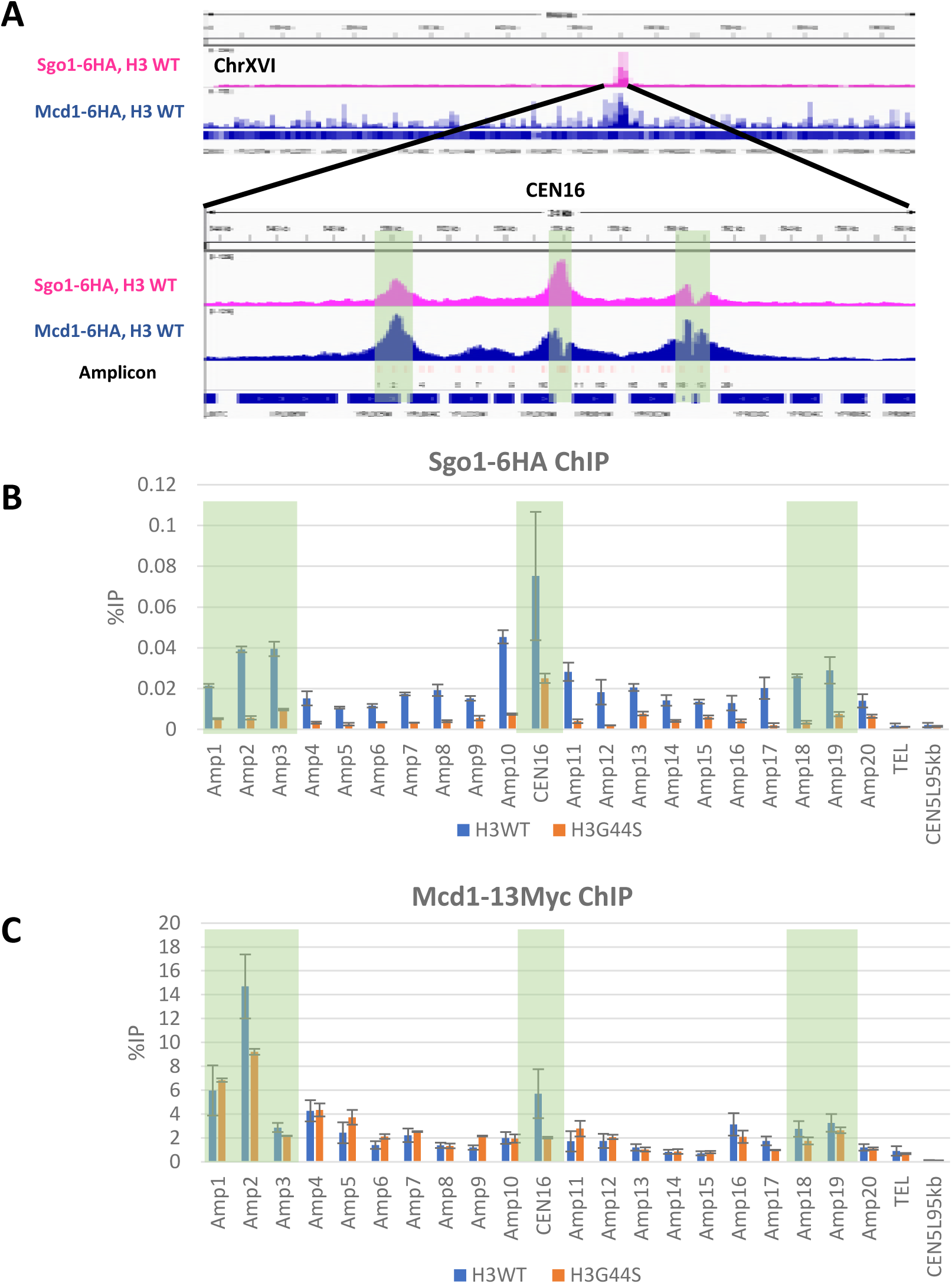
The histone H3 tension sensing motif is essential for pericentric Sgo1p localization but not Mcd1p. A. Distribution of Sgo1p (magenta) and Mcd1p (blue) across chromosome XVI as revealed by ChIP-seq. The centromeric region is blown up to show the detail distribution of these two proteins. PCR amplicons are enumerated and shown in light pink bars below the Mcd1 peaks. The open reading frames and their transcription directions are shown at the bottom. B and C. Quantitative real-time PCR analysis of separate ChIP experiments. Sgo1p-HA and Mcd1p-Myc (both expressed from their native loci) were ChIP’ed from cells bearing the wildtype or a mutant TSM (G44S). The three enrichment sites are marked with the shaded boxes.

**Figure 4.**
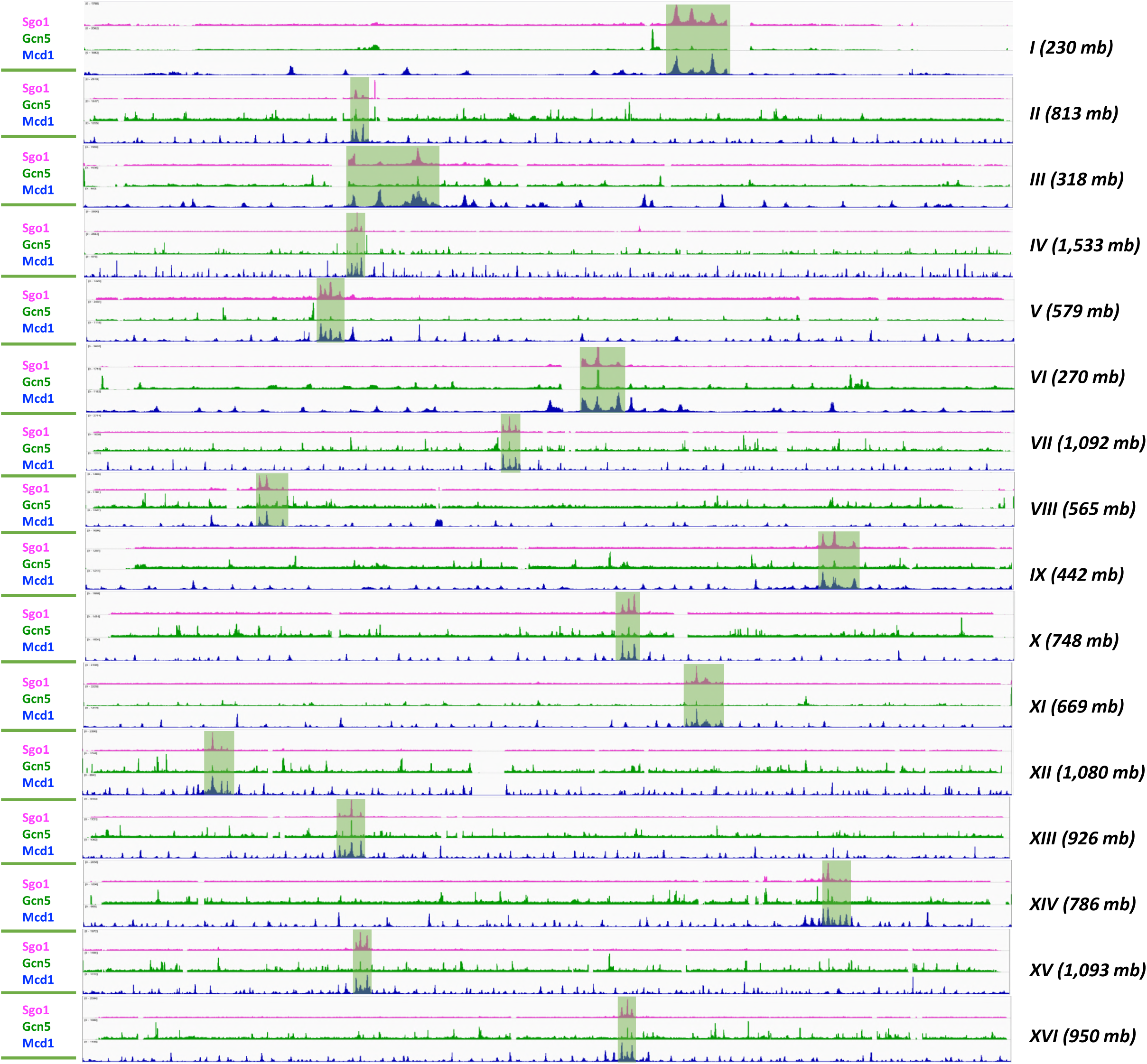
Gcn5p is enriched in centromeres but shows no overlap with cohesin elsewhere. Genome-wide distribution of Gcn5p is compared with that of Sgop1 (magenta) and Mcd1p (blue). The trident Sgo1p localization domain in each chromosome is marked with the shaded boxes.

In addition to comparing our Sgo1p ChIP-seq data with a published Mcd1p dataset (Verzijlbergen et al., 2014), we conducted another set of ChIP assays and used quantitative PCR to examine the localization of Mcd1p and Sgo1p in the same genetic background. To this end, Sgo1p-HA and Mcd1p-Myc expressed from their native loci were subjected to ChIP. DNA products were then examined by quantitative PCR for 21 amplicons that spanned 11 kb of the centromeric region on chromosome XVI, including the three CEN and PC peaks (shaded boxes, Figure 3A top panel). Discrete peaks and valleys are readily visible and show a very high degree of overlapping between Sgo1p and Mcd1p with the ChIP-qPCR data. Additional qPCR analysis of chromosome I amplicons equivalent to those of chromosome XVI also verifies the ChIP-seq observations (Supplemental Figure 2). In addition, parallel ChIP reactions were conducted in the G44S *tsm-* background. While the Sgo1p signals diminish significantly in this region (orange bars, Figure 3C), the Mcd1p-Myc enrichment is not significantly affected, which demonstrates that TSM is required for the retention of Sgo1p, not Mcd1p, at pericentromere.

The exceptional selectivity of Sgo1p for a subset of cohesin localization motifs prompted us to compare its genome-wide distribution to that of Gcn5p in mitotic chromosomes. Gcn5p is a critical transcription regulatory histone acetyltransferase. In mitosis, Gcn5p negatively regulates the tension sensing motif (Luo et al., 2016), and is important for maintaining the normal centromere chromatin structure (Vernarecci et al., 2008). Consistently, Gcn5p is present at mitotic centromeres (Luo et al., 2016). To see whether Gcn5p exhibits a mitotic chromosome localization pattern similar to that of Sgo1p, ChIP-seq was conducted on a Myc-tagged Gcn5p. The results show that, while Gcn5p is found enriched at all centromeres, its pericentric presence is practically negligible (shaded boxes showing CEN/PC peaks of Sgo1p, Figure 3). Importantly, throughout the genome, there is very little overlapping between Gcn5p and Mcd1p enrichment. This is not unexpected for Gcn5p is recruited to the 5’ region of many genes for transcriptional regulation, but Mcd1p and the rest of the cohesin complex are enriched at the intergenic region of convergent genes. There appears to be an enrichment of Gcn5p at RNA polymerase III-controlled targets, such as tRNA genes. These ChIP-seq results are consistent with the canonical roles of Gcn5p in transcription (Venters et al., 2011), although we do not exclude the possibility that at least part of the mitotic distribution pattern of Gcn5p might be for chromatin metabolism during mitosis. Together, ChIP-seq data presented above reveal unique association between Sgo1p and Mcd1p at and near the centromeres. However, this connection does not apply to the recruitment of Gcn5p, indicating a specific functional interplay between Sgo1p and the cohesin complex.

### Pericentric localization of Sgo1p depends on local cohesin enrichment

To better understand the contribution of cohesin to the chromosomal distribution of Sgo1p, we took two approaches. Firstly, we deleted the *IML3* gene that encodes a subunit of the Ctf19 kinetochore subcomplex (Pot et al., 2003, Ghosh et al., 2001). Iml3p is required for pericentric localization of cohesin (Kiburz et al., 2005, Fernius and Marston, 2009). Using an *iml3Δ* strain, we tried to answer whether disrupting pericentric Mcd1p domain would also impair Sgo1p retention. Figures 5A and 5B show that the enrichment of both Mcd1p and Sgo1p is reduced to the background level (i.e., the telomeric region) in cells lacking the *IML3* gene (orange bars), whereas the arm cohesin remains unaltered (Figure 5A, CEN1L100 kb, CEN4R595 kb, and CEN16L465 kb).

**Figure 5.**
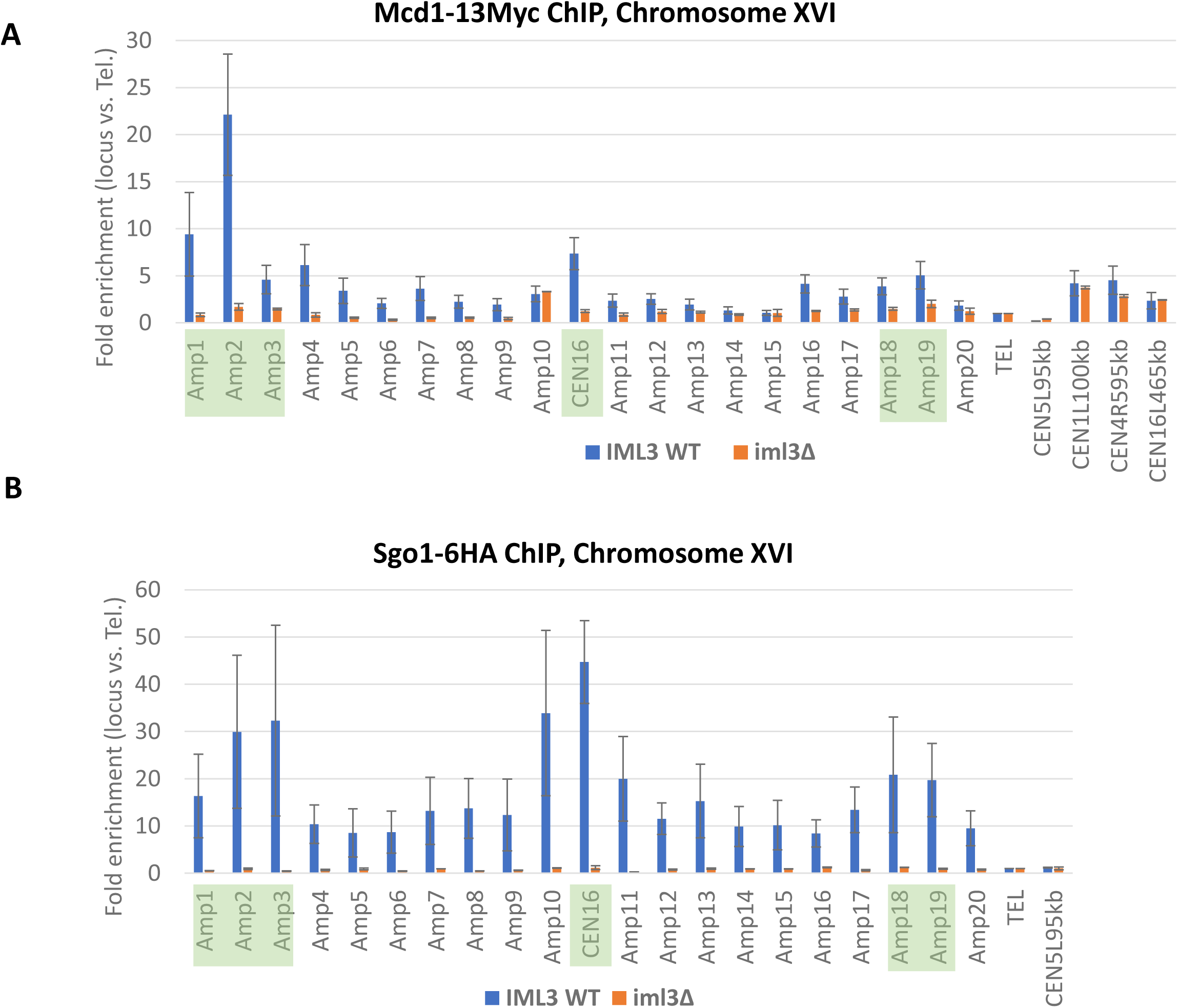
Sgo1p recruitment requires cohesin. *IML3*^*+*^ and *iml3Δ* cells were arrested in G2/M before ChIP for the localization of Mcd1p (panel A) and Sgo1p (panel B). The enrichment of these two proteins was quantified by qPCR. Amplicons are same as those in Figure 3. Shown are average of two biological replicas.

In the second, more specific approach to assessing the importance of cohesin in Sgo1p localization, we targeted a specific cohesin associated region (CAR) on chromosome IV for inducible disruption. Half of all CARs are in the intergenic region of two convergent transcription units (Glynn et al., 2004). Driving transcription through an Mcd1p enrichment site is expected to dislodge both Mcd1p and Sgo1p. To test this prediction, we changed the pericentric CAR between YDR004W and YDR005C to a galactose-inducible promoter *GAL1* (*pGAL1*, Figure 6A). Replacing CAR with *pGAL1* does not affect Mcd1p or Sgo1p if cells were grown on the non-inducing sugar raffinose (blue bars, Figures 6B and C). Galactose addition activated transcription of YDR004W by twofold (Supplemental Figure 3), and also caused Mcd1p signal at amplicons 7 and 8 to diminish (compare orange and blue bars, Figure 6B) whereas the centromeric signal (amplicon 1, 12 kb on the left of *pGAL1*) was unaffected. Similarly, the Sgo1p signal at amplicons 7 and 8, but not 1, was significantly reduced as well. We therefore conclude that active transcription can perturb the establishment of Mcd1p and Sgo1p domain locally. From Figures 3, 5, and 6, we conclude that the pericentric localization of Sgo1p requires both the TSM and the cohesion complex.

**Figure 6.**
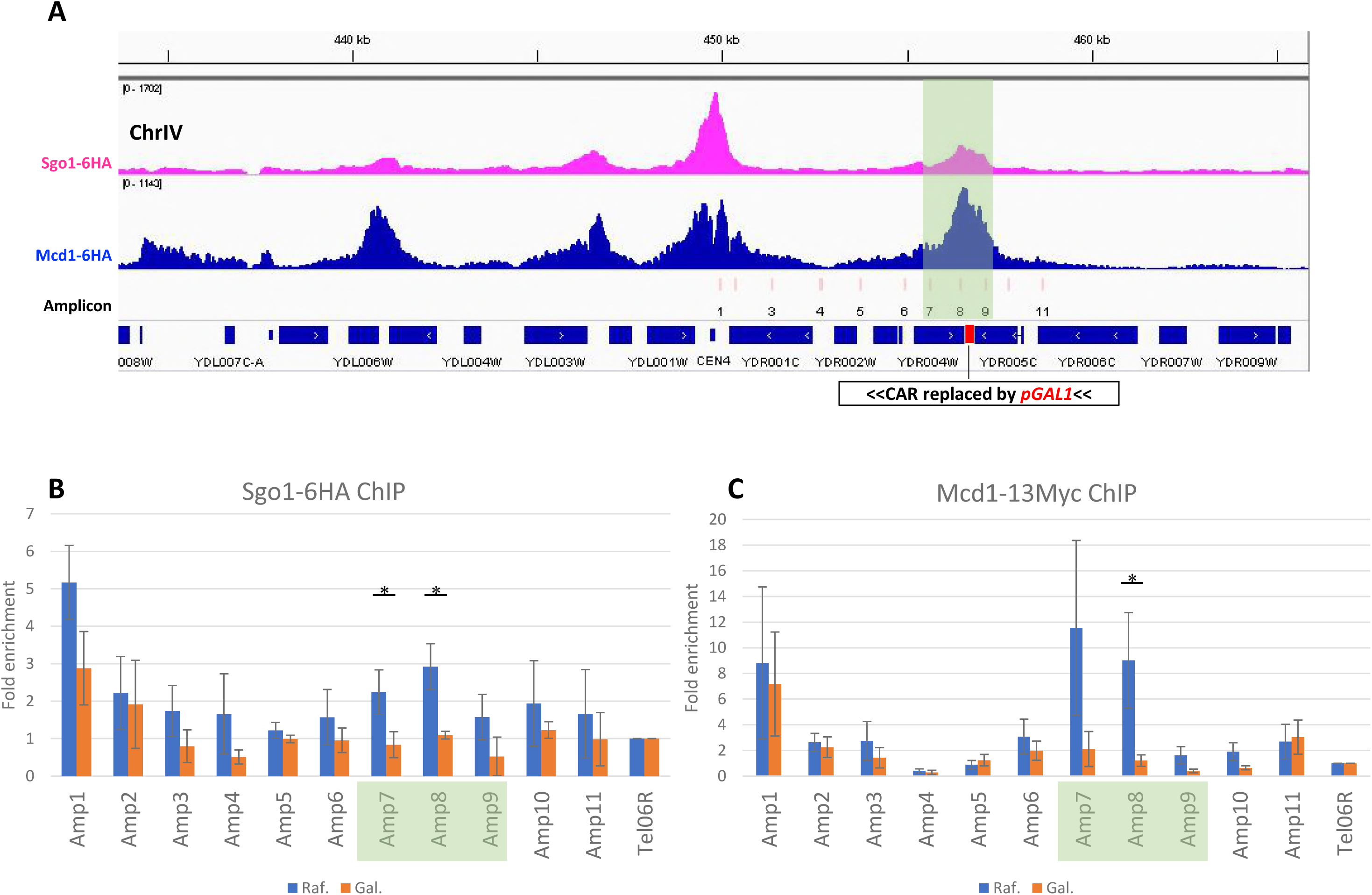
Pericentric localization of Sgo1p and Mcd1p is susceptible to ectopic transcription through the region. A. Blow-up of the *CEN4* area showing the major Sgo1p and Mcd1p peaks, amplicons for quantitative PCR in light pink, and the position of the ectopic *GAL1* promoter (*pGAL1*, in red). *pGAL1* drives galactose-induced transcription toward *yDR004W*. Amplicons 7 – 9 are highlighted by a shaded box. B and C. ChIP and quantitative PCR results of Sgo1p and Mcd1p localization in the absence (Raf., raffinose) or presence (Gal., galactose) of transcription from the *pGAL1* promoter. Fold-enrichment of each amplicon was obtained by comparing with the control corresponding to the subtelomeric region of chromosome VI (*TEL06R*). Error bars are standard deviations from three biological replicas * indicates p-value < 0.05.

### Pericentric Sgo1p domain formation does not involve intervening valley regions

Sgo1p docks on centromeres via direct association with Bub1p-phosphorylated Ser121 of histone H2A (phos.H2A) within the single centromeric nucleosome (Kawashima et al., 2010). Sgo1p also binds the N’ tail of the centromere-specific histone H3 variant, Cse4p (Mishra et al., 2017). It is likely that phos.H2A and Cse4p provide the docking site for Sgo1p that nucleates outward spread toward the pericentric regions. The establishment of PC enrichment of Sgo1p may be accomplished by one of two mechanisms. In the rippling mode, a wave of Sgo1p spreads along the nucleosomal path before it stops and accumulates at the first cohesin block. Alternatively, Sgo1p “leaps” directly from centromeres to the PC region where it is retained by the tension sensing motif. In both modes, Sgo1p is underrepresented at the region between the CEN and PC peaks, resulting in the “valleys” seen in the two-dimensional presentation of the ChIP-seq results. These two modes of Sgo1p recruitment can be differentiated by examining the dynamics of CEN and PC peaks emergence when cells progress through mitosis. An intermediate stage where a significant elevation of Sgo1p signals at the valley region before they move outward to generate the final PC peaks would support the rippling mode. To test these two models, we tagged Sgo1p and Mcd1p in the same strain to avoid any variation between cells with different genotypes. Cells expressing Sgo1-6HA and Mcd1-13Myc were arrested in G1 phase by the pheromone α factor. They were then released into the division cycle before collection at 30, 37.5, 45, 52.5, 60, 75, and 90 minutes after the release. Budding index revealed the timing of the progression through mitosis during the course of experiments (Figure 7A). ChIP results (Figure 7B and Supplemental Figure 4) show that Sgo1p was first detectable at *CEN16* 37.5 minutes after release from G1 arrest, when cells were at the juncture of G1 and S phases. This is also when Sgo1p expression starts (Indjeian et al., 2005). While Sgo1p centromeric abundance continued to rise, the adjacent PC peaks started to surface in the next 7.5 minutes (amplicons 3, 16, and 21). These signals culminated at T_60’_ (green bars, Figure 7B) and diminished afterwards (T_75’_ and T_90’_). Between T_60’_ and T_75’_, approximately 20% of cells entered the anaphase (green sector, Figure 7A), indicating that biorientation had been established in this population of cells. The concomitant reduction of Sgo1p signals is in excellent agreement with the tension-dependent removal of Sgo1p from the chromatin (Nerusheva et al., 2014).

**Figure 7.**
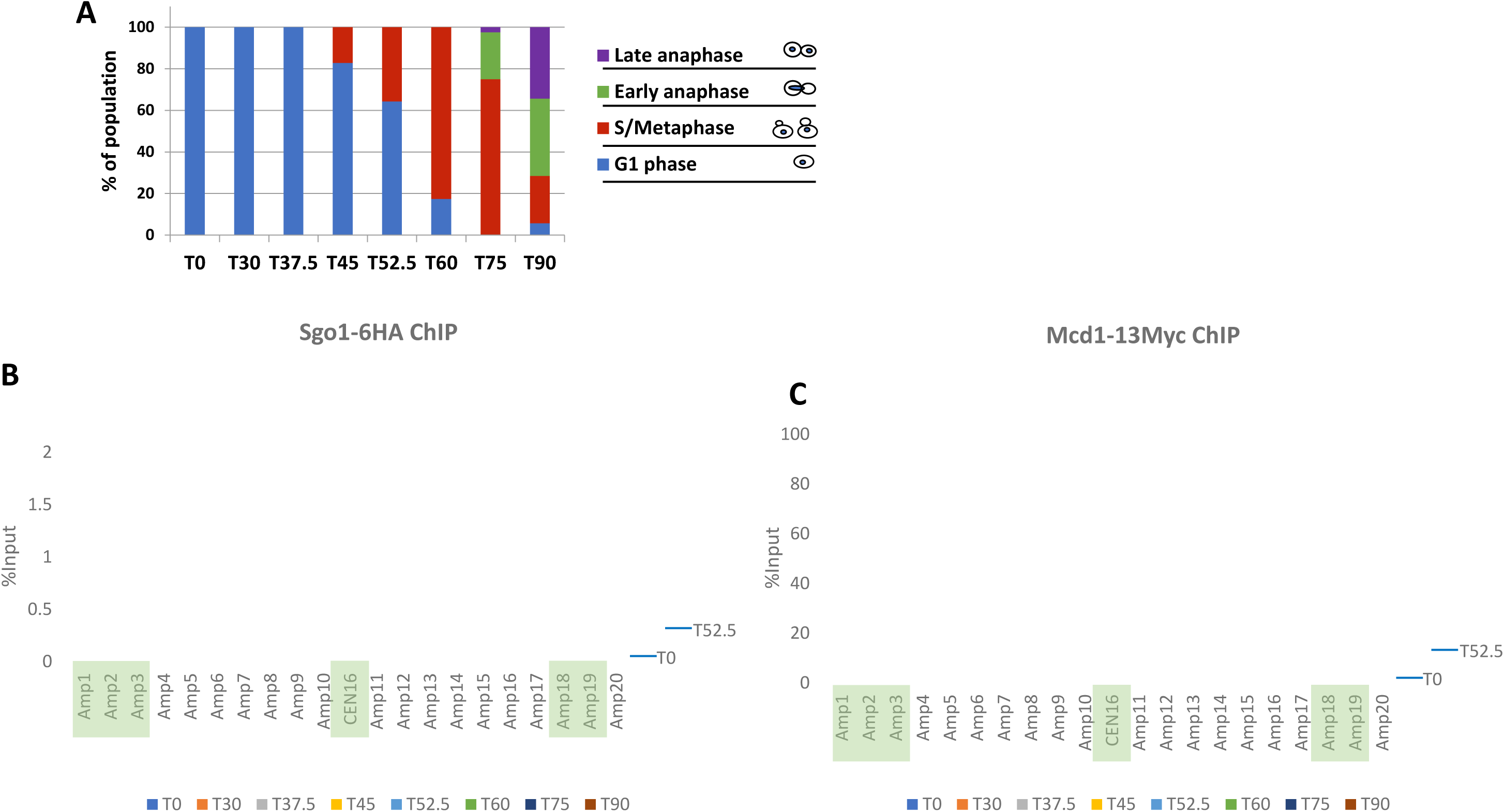
Dynamic recruitment of Sgo1p and cohesin at centromere and pericentromere through cell cycle. A. Budding index of cells collected from the indicated time points. B and C. Sgo1p-HA and Mcd1p-Myc co-expressed in the same cells were examined by ChIP-qPCR. PCR amplicons correspond to *CEN16* and nearby regions. See Figure 3 for positions of these amplicons.

The kinetics of Mcd1p association with CEN and PC exhibited several important distinctions. Firstly, while Mcd1p signals jumped at T30’, the three subsequent time points (T37.5’, T45’ and T52.5’) saw a reduction of the overall Mcd1p signals, which then climbed up again, and peaked at T75’ before disappearance by T90’, when the majority of cells passed the metaphase-to-anaphase transition (Figure 7A). The dynamic changes before T60’ probably resulted from transcriptional activities in S and G2 phases. The abrupt increase of Mcd1p signal at T60’ agreed well with the budding index that 80% of the cells were in the metaphase when cohesion of sister chromatids was most critical. Lastly, the highest levels of the Mcd1p abundance were found to be at T75’ before its quick disappearance by T90’, both were 15’ later than Sgo1p. The different kinetics of Sgo1p and Mcd1p dissolution concurs with the anticipated sequence of biorientation, Sgo1p removal, and Mcd1p cleavage that marks anaphase onset.

One critical observation from results in Figure 7 is that during the formation of the Sgo1p CEN and PC tripartite motif, the two valleys flanking the CEN peak never rose to the levels of PC at any given time. That the PC peak-to-peak distance persists throughout their lifespan in mitosis argues strongly against the notion that Sgo1p spreads along consecutive nucleosomes from CEN to PC. Rather, these data support the model that Sgo1p either “hops” from CEN to PC, or is recruited simultaneously to these regions to generate the tripartite motif.

### Chromosome conformation capture reveals correlation between Sgo1p enrichment and chromatin architecture

If Sgo1p targets its pericentric destination immediately after or concomitantly with the centromeric recruitment, it seems likely that the PC regions are rendered accessible to Sgo1p whereas the intervening regions are somehow hidden from Sgo1p. Because the interaction between Sgo1p and TSM does not require any posttranslational modification (Luo et al., 2016, Luo et al., 2010), a non-epigenetic feature may distinguish the PC Sgo1p targets from other areas near the centromeres. We felt that chromatin architecture would be a good candidate that dictates the (in)accessibility of the CEN/PC region to Sgo1p. Compaction of chromatin in mitosis involves condensin and cohesin complexes (Hudson et al., 2009, Mehta et al., 2013). Both complexes are also shown to be critical for organizing pericentromere in prometaphase (Stephens et al., 2011, Yeh et al., 2008, Nasmyth, 2011). Cohesin facilitates the formation of intrachromosomal centromeric loops for mitotic segregation and resides near the summits of these loops. On the other hand, the condensin complex holds and organizes the bottom of these loops along the spindle axis (Stephens et al., 2011). Taking together these models and our results shown above, we suspect that higher-ordered chromosomal architecture, e.g., chromosome looping, might be part of the mechanism underlining the highly selective pericentric localization for Sgo1p.

If Sgo1p recruitment is linked to chromosome looping in mitosis, we predicted that PC and CEN peaks of Sgo1p were spatially near each other owing to the action of such proteins as cohesin and condensin. This hypothesis was tested by chromosome conformation capture (3C) (Dekker et al., 2002). Yeast nuclei were harvested from G1 and G2/M arrest and were subjected to *Eco*R I digestion with or without formaldehyde fixation, followed by ligation under a condition that favored intramolecular ligation. The resultant DNA libraries were analyzed by PCR using one of two centromere-proximal anchor primers, oXD159 for *CEN1* and oXD162 for *CEN16*. In each quantitative PCR reaction, these anchor primers were paired with a distal primer that is 3 – 50 kb away (black arrows, Figure 8A). All primers hybridized to the same strand of DNA, hence should not produce any PCR product without the 3C treatment. On the other hand, ligation at the anticipated *Eco*R I sites after formaldehyde fixation would generate templates amplifiable by the anchor and the locus-specific primers. Comparing the intensity of PCR products amplified from samples with or without formaldehyde treatment yielded “crosslinking frequency” that is indicative of the propensity for the two primer target regions to be spatially brought together by chromatin-associating factors.

**Figure 8.**
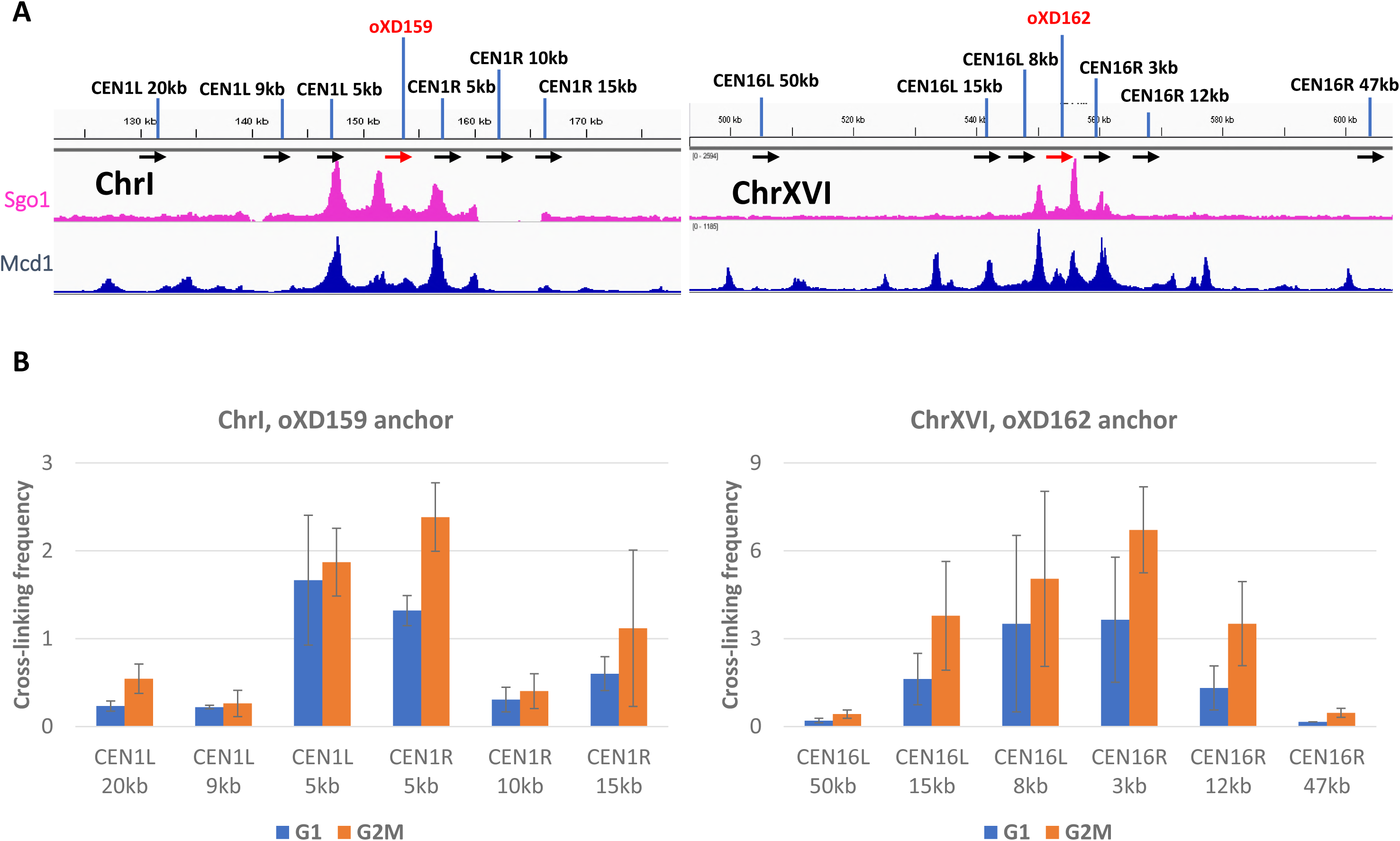
Sgo1p tripartite localization domain is associated with high-ordered chromatin architecture in mitosis. Chromosome conformation capture (3C) assay was used to examine chromatin looping near *CEN1* and *CEN16*. Cells arrested in G1 or G2/M phase were fixed with formaldehyde, and the isolated nuclei treated with *Eco* RI before DNA ligation. An identical amount of final ligated DNA library was amplified by PCR using one of two common anchor primers (oXD159 and oXD162 for chromosomes I and XVI, respectively; red arrows) against different locus-specific primers (black arrows; named for their distance to the centromere, L = left; R = right) 3 – 50 kb away. All primers face toward the same direction. PCR products were resolved by gel electrophoresis and quantified with the NIH Image J software. Shown are the signals relative to the same amplicons without formaldehyde crosslinking. Error bars are standard deviations from three biological replicas.

The 3C assays indeed show that, after crosslinking, the centromeric primers oXD159 and oXD162 could amplify with primers hybridizing to Mcd1p peaks that were 3 to 15 kb away (e.g., oXD159 + CEN1L 5kb or CEN1R 5kb, and oXD162 + CEN16L 8kb or CEN16R 3kb; Figure 8B). Some of the amplification products spanned a region with a conspicuous Mcd1p signal without Sgo1p (e.g., oXD159 + CEN1L 20kb, oXD162 + CEN16L 15kb), consistent with the idea that chromosomal loops generated by the cohesin complex is a prerequisite for Sgo1p localization (Stephens et al., 2011, Verzijlbergen et al., 2014). The crosslinking frequency from G2/M nuclei was in general higher than G1 (orange vs. blue bars), which indicates that the nuclear architecture climaxes during mitosis, but may be partially preserved after exiting from M phase. This notion is consistent with the weak but readily recognizable Mcd1p peaks in cells arrested at G1 (Figure 7).

## DISCUSSION

This work depicts the genome-wide localization of Sgo1p in mitotic *S. cerevisiae* cells. On each chromosome, Sgo1p displays a tripartite localization domain consisting of a middle centromeric and typically two flanking pericentric peaks. Sgo1p co-localizes with the cohesin complex. However, despite that cohesin is recruited to numerous loci across the genome, Sgo1p only rendezvouses with the centromeric and the adjacent pericentric cohesin. This confined localization of Sgo1p requires an intact tension sensing motif of histone H3. Ectopic transcription that disrupts pericentric cohesin localization also dislodges Sgo1p in situ. Overexpression causes Sgo1p to expand its presence, but the new Sgo1p peaks have high propensity to co-localize with cohesin. This unique trident shape of Sgo1p domain on each chromosome appears to be associated with chromatin looping in mitosis, thus linking higher-ordered chromatin architecture to positioning Sgo1p for the crucial tension sensing function of segregation.

Studies of yeast and human cells have demonstrated the importance of cohesin in Sgo1p recruitment to pericentromere (Kiburz et al., 2005, Liu et al., 2013a). However, cohesin alone is not sufficient for the pericentric retention of Sgo1p. The tension sensing motif of H3 is also required for keeping Sgo1p in this region to ensure error-free segregation. While a Gly-to-Ser mutation in the TSM has no effect on cohesin localization, both pericentric and, to a lesser extent, centromeric enrichment of Sgo1p is compromised ((Luo et al., 2010) and Figure 3C). The establishment of the centromeric and pericentric domain of Sgo1p likely follows a spillover model in that Sgo1p is first recruited to the centromeres via direct association with Cse4p (Mishra et al., 2017) and histone H2A phosphorylated at Ser121 by kinase Bub1p (Kawashima et al., 2010, Fernius and Hardwick, 2007). Congregation of Sgo1p molecules at centromeres permits its spread to the adjacent pericentric nucleosomes where cohesin has already been loaded. This spread may result from the turnover of the association between Sgo1p and centromeric proteins. Alternatively, the homodimerization activity of Sgo1p, evidenced by yeast two-hybrid tests (Mishra et al., 2017), may facilitate the growth of the Sgo1p domain from centromeres to pericentric regions where the cohesin complex resides. By binding to nucleosomes, cohesin may also help to make the tension sensing motif more accessible for Sgo1p before biorientation is established (Fernius and Hardwick, 2007, Kawashima et al., 2010, Luo et al., 2010, Luo et al., 2016). Due possibly to the total pool size of Sgo1p, it only spreads to the first and nearest cohesin cluster. Overexpression of Sgo1p can further its spread primarily to adjacent pre-existing cohesin conglomerates (Figure 2).

The distinct kinetics of engaging Sgo1p and cohesin (Mcd1p) at *CEN16* (Figure 5) and *CEN1* (Supplemental Figure 4) is consistent with the notion that cohesin organizes chromatin into a platform for mitotic machinery to execute error-free segregation. Mcd1p appears earlier than Sgo1p does, but fluctuates in abundance before metaphase. In the meantime, Sgo1p continues to accumulate at CEN and PC peaks until it reaches the maximum. When cells enter anaphase, Sgo1p dissipates. It is critical that before Mcd1p levels climb to the highest, Sgo1p already starts disappearing from CEN and PC regions (compare T_60’_ and T_75’_, Figure 5 and Supplemental Figure 4). This time difference echoes the report of tension-dependent removal of Sgo1p from chromatin at the juncture of metaphase and anaphase (Nerusheva et al., 2014), and is consistent with the model that the removal of Sgo1p from chromatin is registered by cells as achieving biorientation.

The centromeric and pericentric clusters of Sgo1p appear almost simultaneously, leaving the intervening regions low in Sgo1p abundance throughout the lifespan of these peaks. Given that the histone H3 tension sensing motif decorates the whole genome and functions without a post-translational modification, the non-continuous nature of the confined Sgo1p peaks on each chromosome strongly suggests physical hindrance in these Sgo1p-free valleys. Our recent findings that Gcn5p acts as a negative regulator for tension sensing motif and Sgo1p functional interaction (Buehl et al., 2018, Luo et al., 2016) alludes to an intriguing possibility that Gcn5p, acetylated H3, or a downstream effector may prevent Sgo1p from binding to the chromosome arms. ChIP-seq data show a lack of correlation between Gcn5p and these Sgo1p-free valleys in mitosis (Figure 3), arguing against a direct, physical role of Gcn5p. Rather, we favor the possibility that a structural feature dictates the accessibility of pericentric chromatin to Sgo1p. Indeed, the chromosome conformation capture results (Figure 8) show that DNA around the centromere loops into a higher-ordered structure that includes centromere and the adjacent Sgo1p and cohesin clusters, a scenario reminiscent of the C-loop model put forth by Bloom and colleagues (Yeh et al., 2008, Salmon and Bloom, 2017). The C-loop conformation posits that pericentric chromatin harbors alternating cohesin and condensin complex clusters. Condensin and the associated chromatin in pericentromeres are restricted to the microtubule axis between spindle pole bodies, whereas cohesin and the cognate CARs are radially positioned, forming the wall of a barrel. In this model, multiple layers of chromatin loops distribute axially, with the top and bottom of this barrel being the clustered centromeres from all 16 chromosomes. Poleward pulling from biorientation stretches the length of this barrel and narrows its diameter.

How does Sgo1p fit into the tension sensing function? Taking together the ChIP-seq and 3C results, we suggest that cohesin is responsible for creating and joining multiple loops in pericentromeres. With centromeres clustering in the center (Jin et al., 1998), these cohesin-capped loops (Figure 9A) can be viewed as a series of concentric circles (Figure 9B). Sgo1p is recruited to the centromere cluster, from which it encroaches radially to the first pericentric cohesin circle (red circles, Figure 9B). Biorientation instigates both intra- and inter-chromosomal tension (Salmon and Bloom, 2017). The increased space between individual nucleosomes causes a conformational change of the tension sensing motif (Luo et al., 2016, Luo et al., 2010) or even nucleosome dissociation from pericentromeres (Lawrimore et al., 2015). In either case, Sgo1p loses its footings and dissipates from chromatin (Figure 9B, green circles). Tension-induced clearance of Sgo1p in pericentromeres signals biorientation to the spindle assembly checkpoint (Nerusheva et al., 2014). Anaphase thus ensues. This model provides a mechanistic explanation for the mitotic delay caused by Sgo1p overexpression (Clift et al., 2009). Biochemical fractionation experiments demonstrated that yeast cells do not to have a soluble pool of Sgo1p, but rather keep all Sgo1p molecules in the CEN/PC region (Buehl et al., 2018). If true, the overall size of the Sgo1p motif on chromosomes (red circles, Figure 9B) would be dictated by the number of Sgo1p molecules. Overexpression raises Sgo1p levels and expands the range of Sgo1p occupancy to the next cohesin circle farther away from the centromere cluster. Consequently, axial separation of kinetochores has to be extended in order to evict the outermost Sgo1p peaks. Assuming that the quantitative removal of Sgo1p from centromeric and pericentric regions signals biorientation, Sgo1p overdose would require more time to clear Sgo1p before anaphase onset, resulting in mitotic delay. On the contrary, deleting Sgo1p or preventing the formation of the pericentric Sgo1p domain by mutating the tension sensing motif would be interpreted erroneously as biorientation, thus triggering precocious anaphase onset and aneuploidy (Luo et al., 2010, Indjeian et al., 2005).

**Figure 9.**
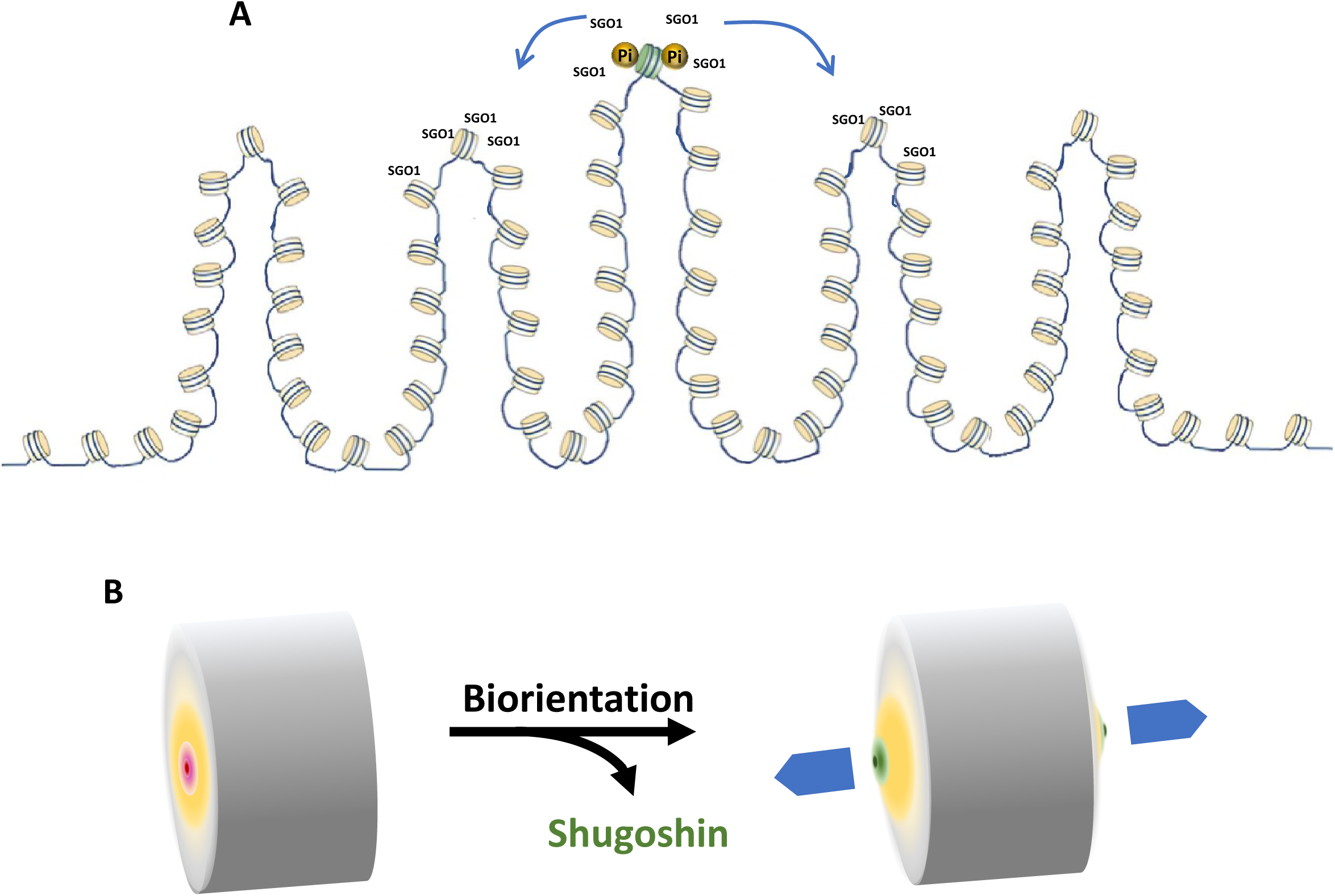
Model for the formation and dynamics of Sgo1p chromatin domain. A. Sgo1p is first recruited to the centromeres via association with phosphorylated histone H2A (Pi). Centromere-bound Sgo1p then spreads to the nearby cohesin-occupied region. B. At the whole genome level, congregation of centromeres aligns the adjacent chromatin loops to form concentric rings (gradient yellow circle) that become the two terminals of the chromatin column. Prior to biorientation, Sgo1p (gradient red circle) resides on the centromere cluster and the first ring of chromatin loops. Poleward pulling from bipolar attachment stretches the centromeric and pericentric chromatin, resulting in a conformational change (gradient green circle) and evicting Sgo1p.

## MATERIALS AND METHODS

### Yeast strains and plasmid constructs

The yeast strains, plasmids, and primers used in this work are listed in Tables 1 to 3. To study the genome wide localization of Sgo1p, the 6HA epitope-tagged Sgo1p strains, yJL345 (H3WT) and yJL346 (H3G44S) were constructed as previous described (Luo et al., 2010). The Sgo1p overexpression strains, yJL322 (H3WT) and yJL324 (H3G44S) were generated by transforming pJL51 (a *URA3* plasmid with p*ADH1*-3HA-*SGO1*-t*ADH1*) into yMK1361 and yJL170, whose endogenous *SGO1* gene was deleted using *TRP1* marker. To ChIP Mcd1p, a 13Myc tag was introduced to the C terminus of *MCD1* locus in yJL347 using pFA6a-13Myc-His3MX6 plasmid as described (Petracek and Longtine, 2002). The resultant strain yXD225 was transformed with either pMK439H3WT or pMK439H3G44S (a *LEU2* plasmid bearing all four core histone genes) and followed by 5-FOA selection to select against pMK440 (a *URA3* plasmid bearing all four core histone genes) containing cells, generating yXD233 (H3WT) and yXD234 (H3G44S). *BAR1* was deleted in yXD233 and yXD234 to yield yXD237 and yXD238 respectively, using homologous recombination approach with *URA3* marker. Another version of *bar1* deletion was made in yXD233 to yield yXD282, using *URA3* recycling approach as described previously (Akada et al., 2002). An adapted *URA3* recycling method was used to replace the CAR sequence between *RAD57* and *MAF1* with *GAL1* promoter. There were 4 steps PCR to attain the recombinant fragment. Step 1, primers oXD236 and oXD237 were used to amplify 3’ end of *RAD57* from genomic DNA. Step 2, amplified *pGAL1* from plasmid pFA6a-*TRP1-pGAL1-3HA* with primers oXD252 and oXD253. Step 3, PCR the *URA3* from plasmid pMK440 using primers oXD254, oXD255 and oXD240. Step 4, combined PCR products from the previous three steps and used primers oXD236 and oXD240 to amplify the final fragment. The resultant DNA was transformed into yXD282 to attain Ura^+^ transformant, which was then subjected to 5-FOA selection to generate yXD286.

**Table 1:**
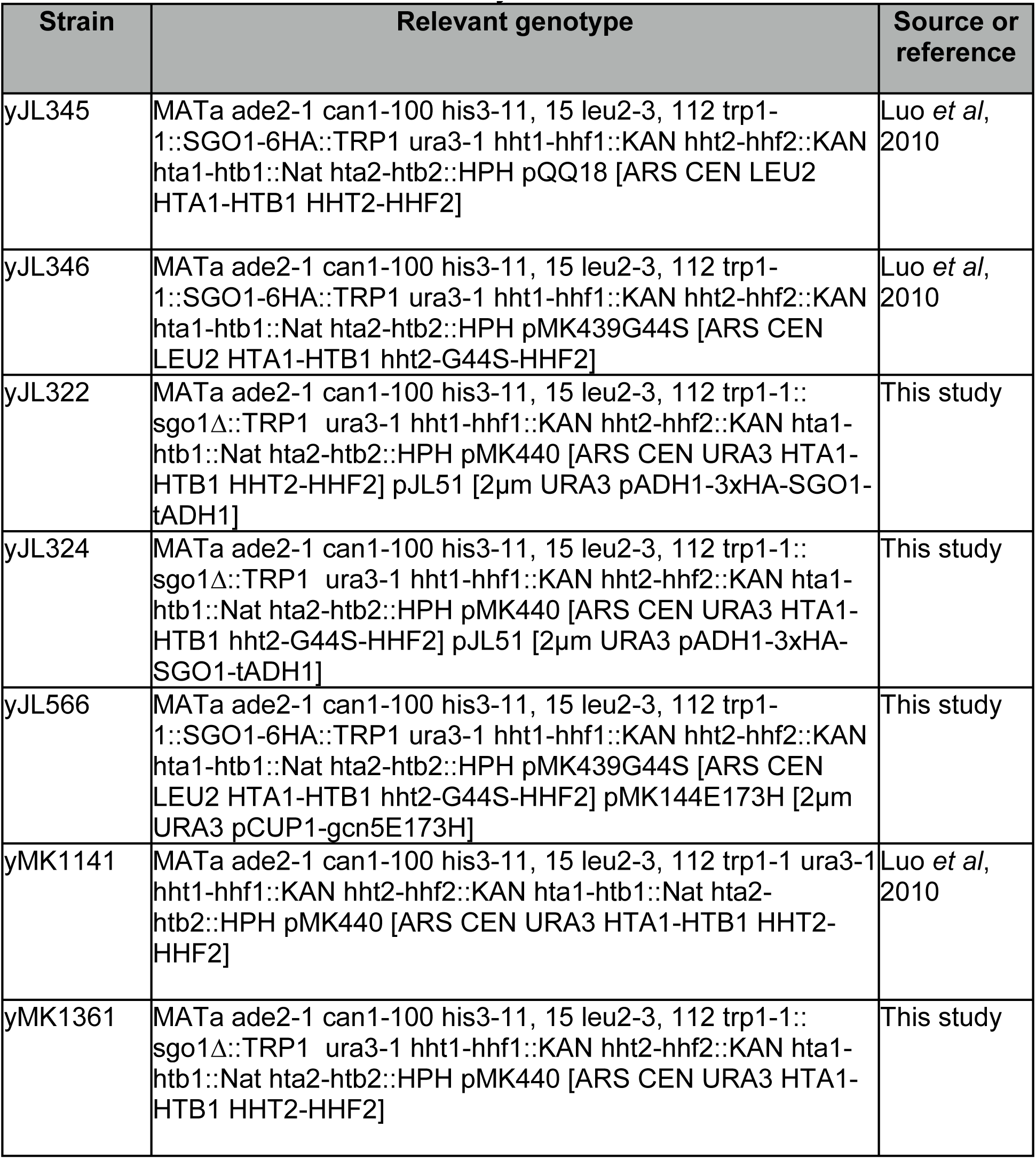
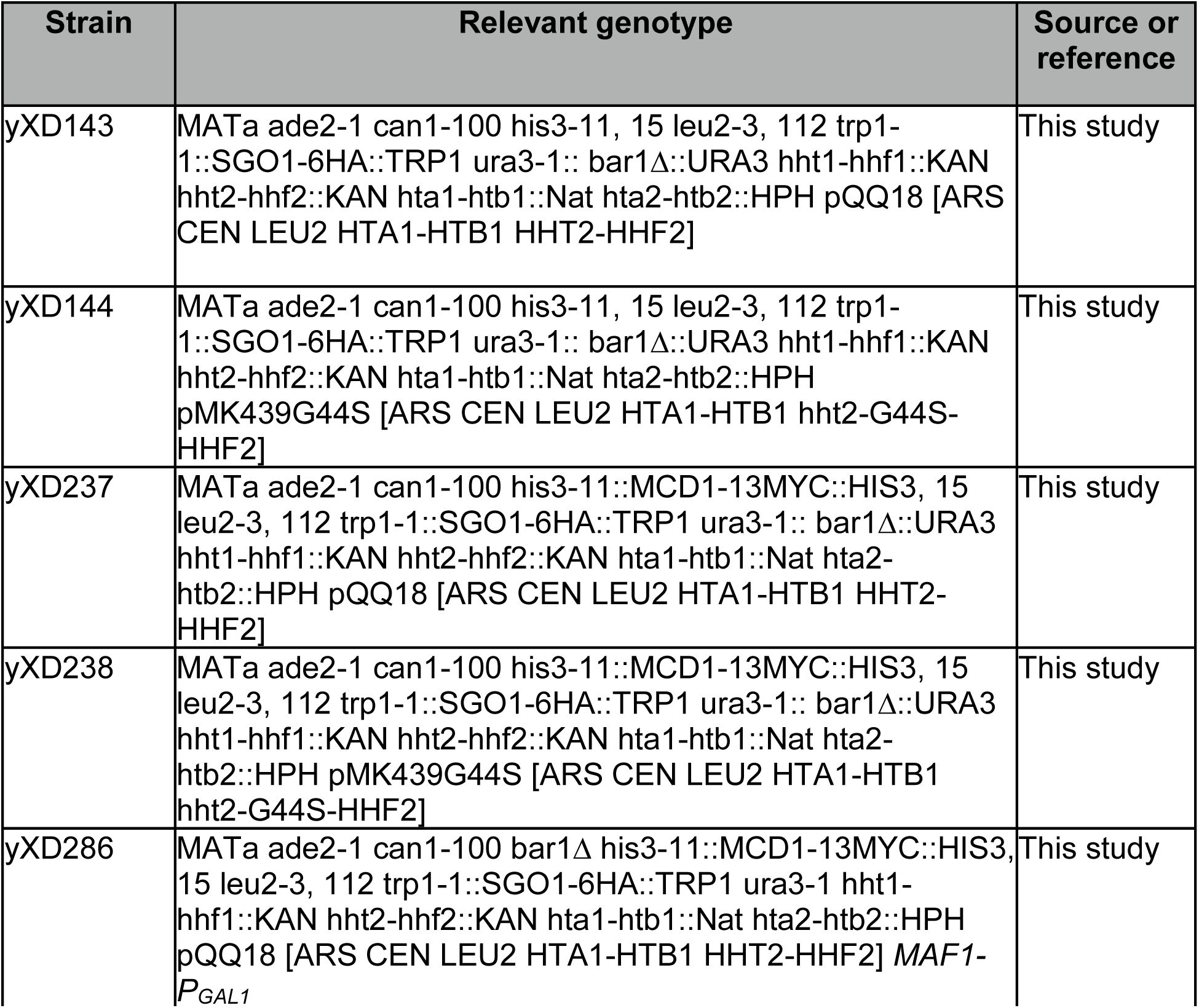
Yeast strains used in this study

**Table 2:**
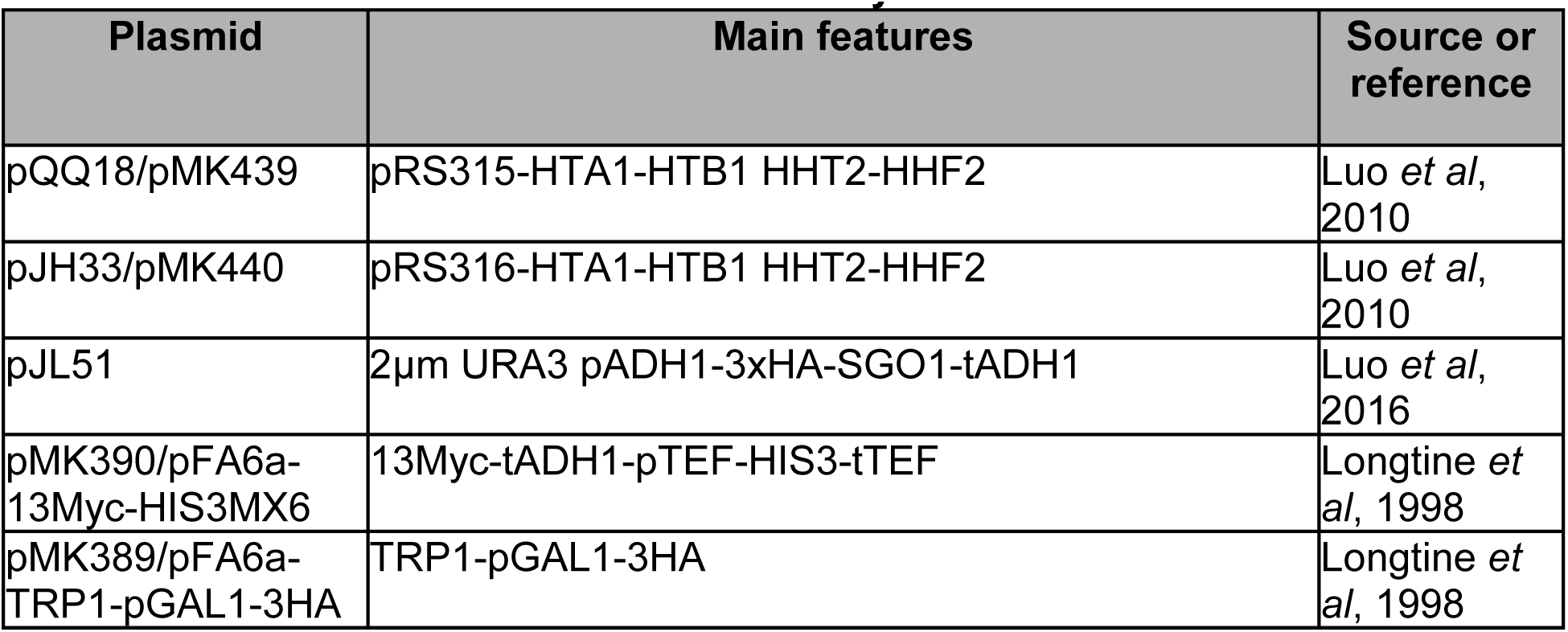
Plasmid constructs used in this study

| Plasmid | Main features | Source or reference |
| --- | --- | --- |
| pQQ18/pMK439 | pRS315-HTA1-HTB1 HHT2-HHF2 | Luo *et al*, 2010 |
| pJH33/pMK440 | pRS316-HTA1-HTB1 HHT2-HHF2 | Luo *et al*, 2010 |
| pJL51 | 2μm URA3 pADH1-3xHA-SGO1-tADH1 | Luo *et al*, 2016 |
| pMK390/pFA6a-13Myc-HIS3MX6 | 13Myc-tADH1-pTEF-HIS3-tTEF | Longtine *et al*, 1998 |
| pMK389/pFA6a-TRP1-pGAL1-3HA | TRP1-pGAL1-3HA | Longtine *et al*, 1998 |

### Yeast methods

Yeast growth media, conditions, and transformation were based on standard procedures (Sherman, 1991). When appropriate, 5% casamino acids (CAA) were used to substitute for synthetic amino acid mixtures as selective medium for uracil, tryptophan, or adenine prototroph. Yeast transformation was done with the lithium acetate method (Gietz et al., 1992).

### ChIP-qPCR and ChIP-seq

ChIP was conducted as previously described (Luo et al., 2010, Kuo and Allis, 1999). To quantify the ChIP results, ChIP DNAs were analyzed with quantitative PCR using primers from Table 3. The libraries of Sgo1p ChIP-seq were prepared as described previously (Ford et al., 2014). 10 ng of ChIP DNA was used for each library preparation. Size selection of libraries was 300-500 bp. Libraries passed quality control were then subjected to Illumina HiSeq 2500 to get 50 bp single-end reads. Reads were mapped to *S. cerevisiae* genome (Saccer 3.0) by Bowtie2 (version 2.2.6) using -m 1 setting for unique matching reads. BEDgraph files of each ChIP-seq experiments were generated by HOMER (version 4.7.2) and were visualized by Intergrative Genomics Viewer (Broad Institute). Read analysis across centromeres was done by using code of Cen-boxplot_100kb.pl adapted from Verzijlbergen et al. (2014). All ChIP-seq data in this study are available at the Gene Expression Omnibus with accession number GSE110953.

**Table 3:**
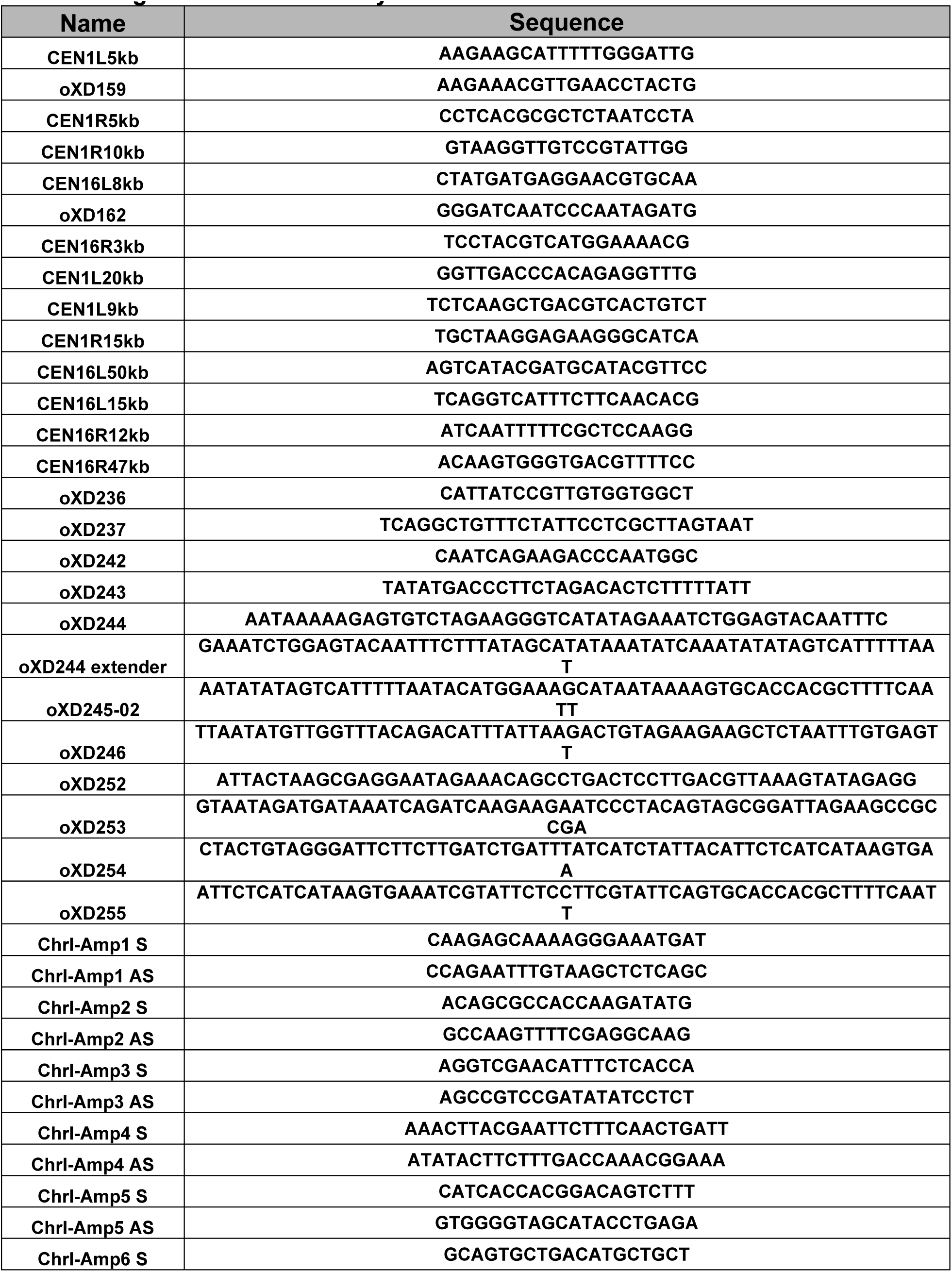
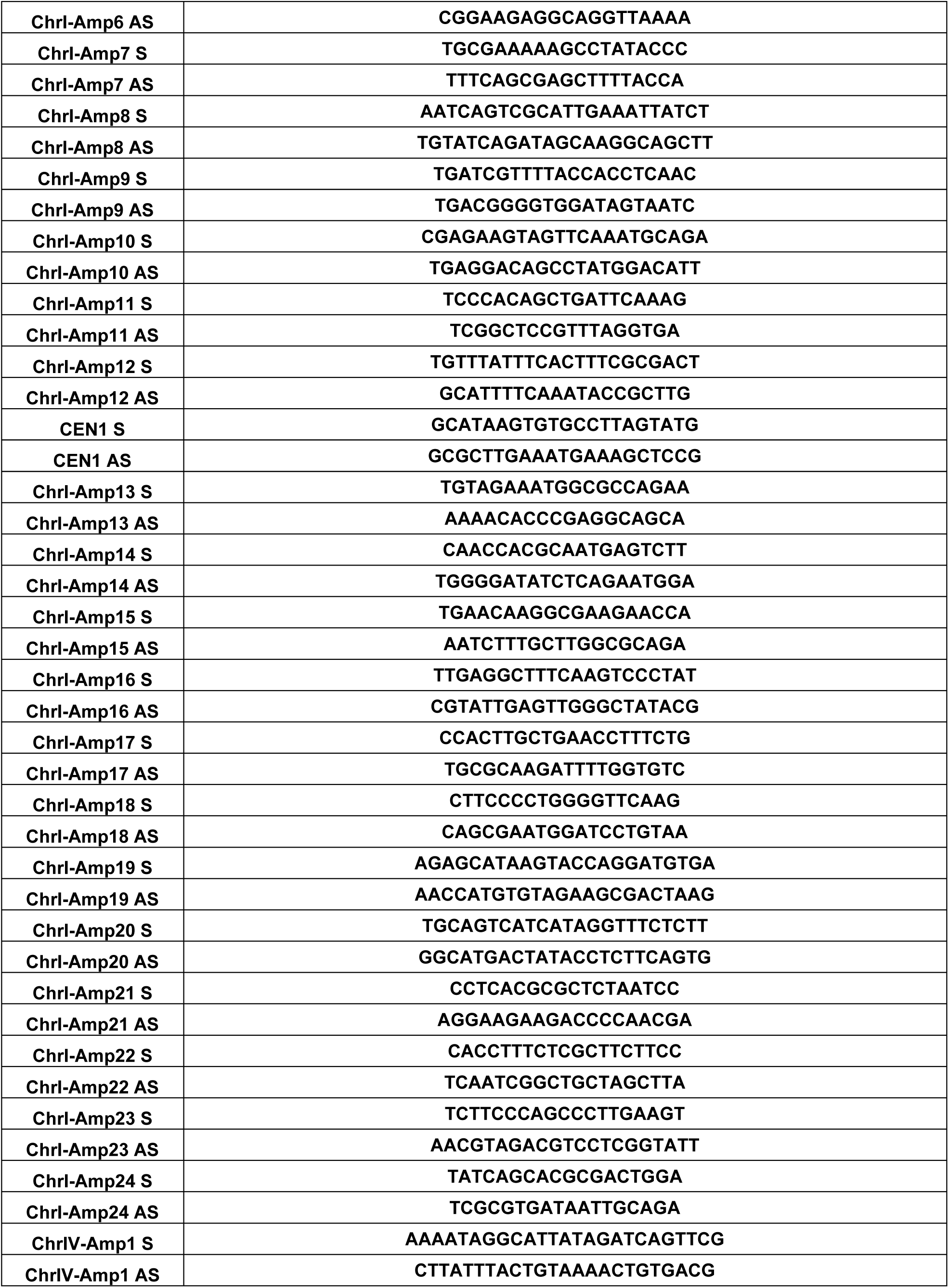
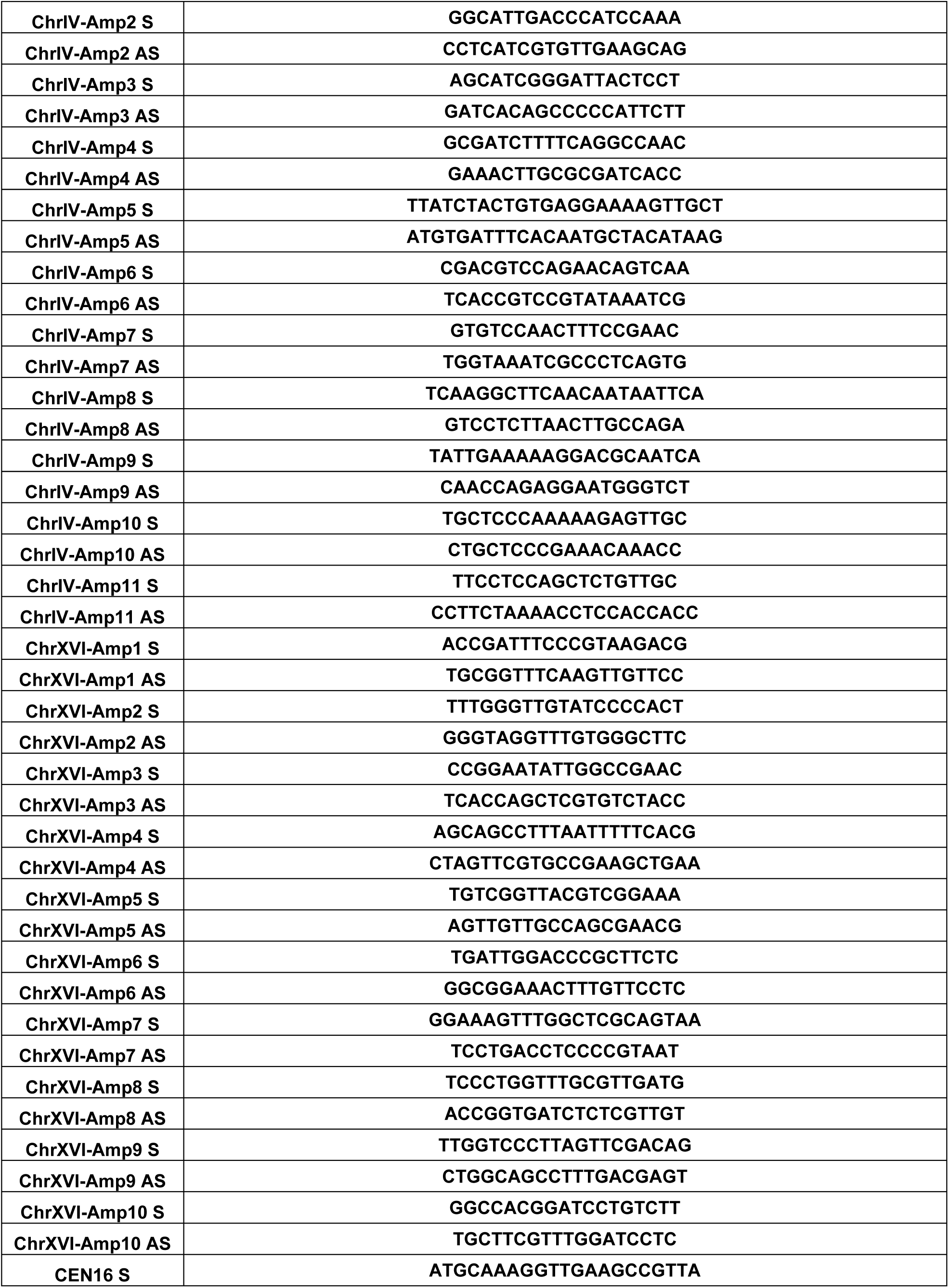
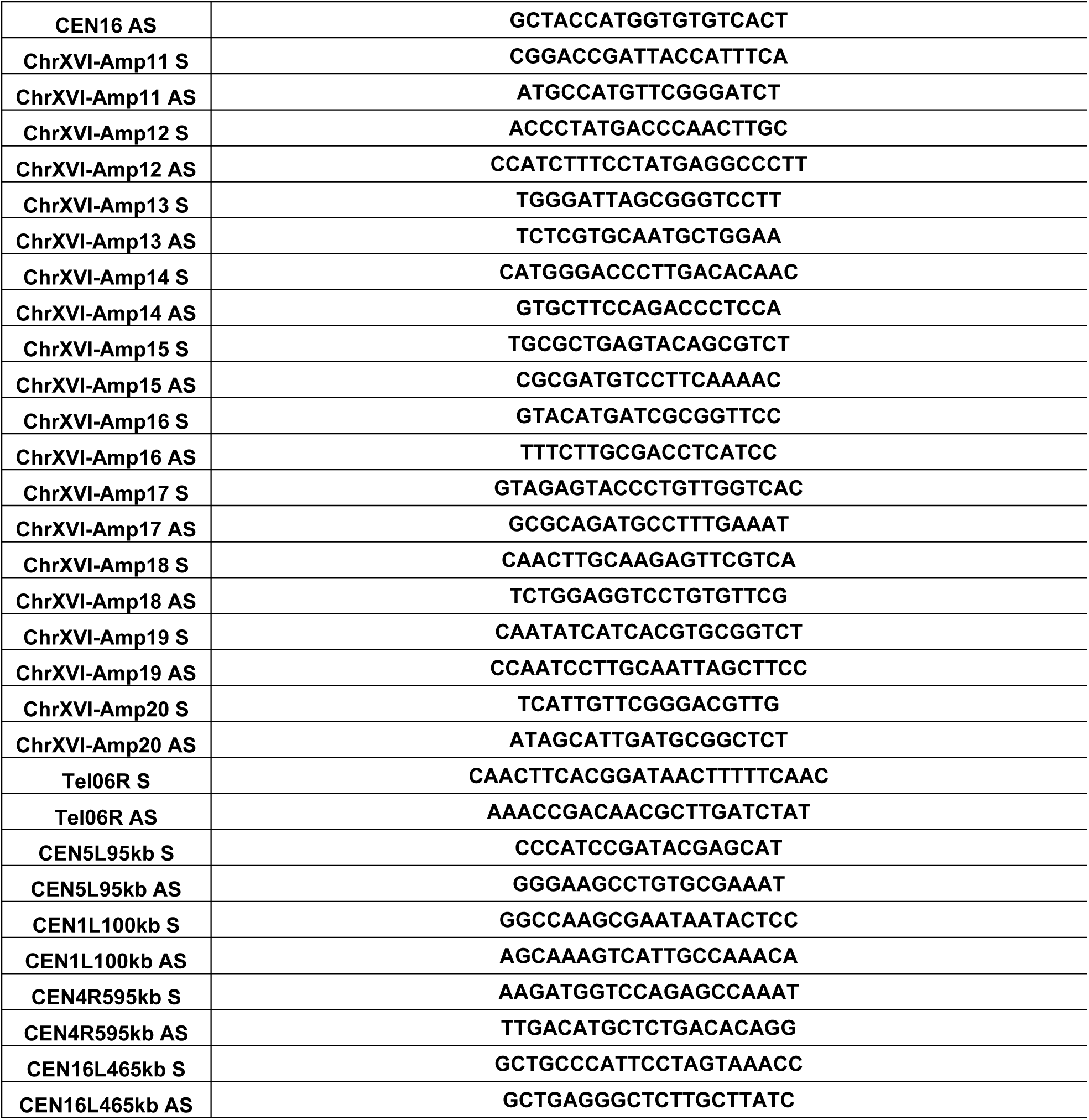
Oligos used in this study

### Chromosome Conformation Capture, 3C

3C was performed in 100 OD_600_ cells of G1 or G2M arrest cells as previously described (Belton and Dekker, 2015). Instead of using mortar and pestle to lyses cells, 50 U/mL lyticase was used to digest the cell wall for 25 min at room temperature. Primers are designed around 50 bp upstream of the targeted *Eco*R I sites. The digestion efficiency of each libraries was evaluated by qPCR. Samples with at least 70% digestion were carried on for following assay. PCR products were resolved by 9% PAGE and stained by ethidium bromide. The intensity of band was analyzed by NIH ImageJ.

## ACKNOWLEDGMENTS

We are grateful for the technical assistance from Monique Floer, Alison Gjidoda, Mohita Tagore, Michael McAndrew, Kurtus Kok, and Sandhya Payankaulam. We thank Christopher Buehl for his critical reading of this manuscript and frequent discussion of the project. This work was supported the National Science Foundation (MCB1050132) and partly by National Institutes of Health (AG051820) to M.-H. Kuo. XD was also supported by a Thesis Completion Scholarship by the Michigan State University.

---

**Supplemental Figure 1.**
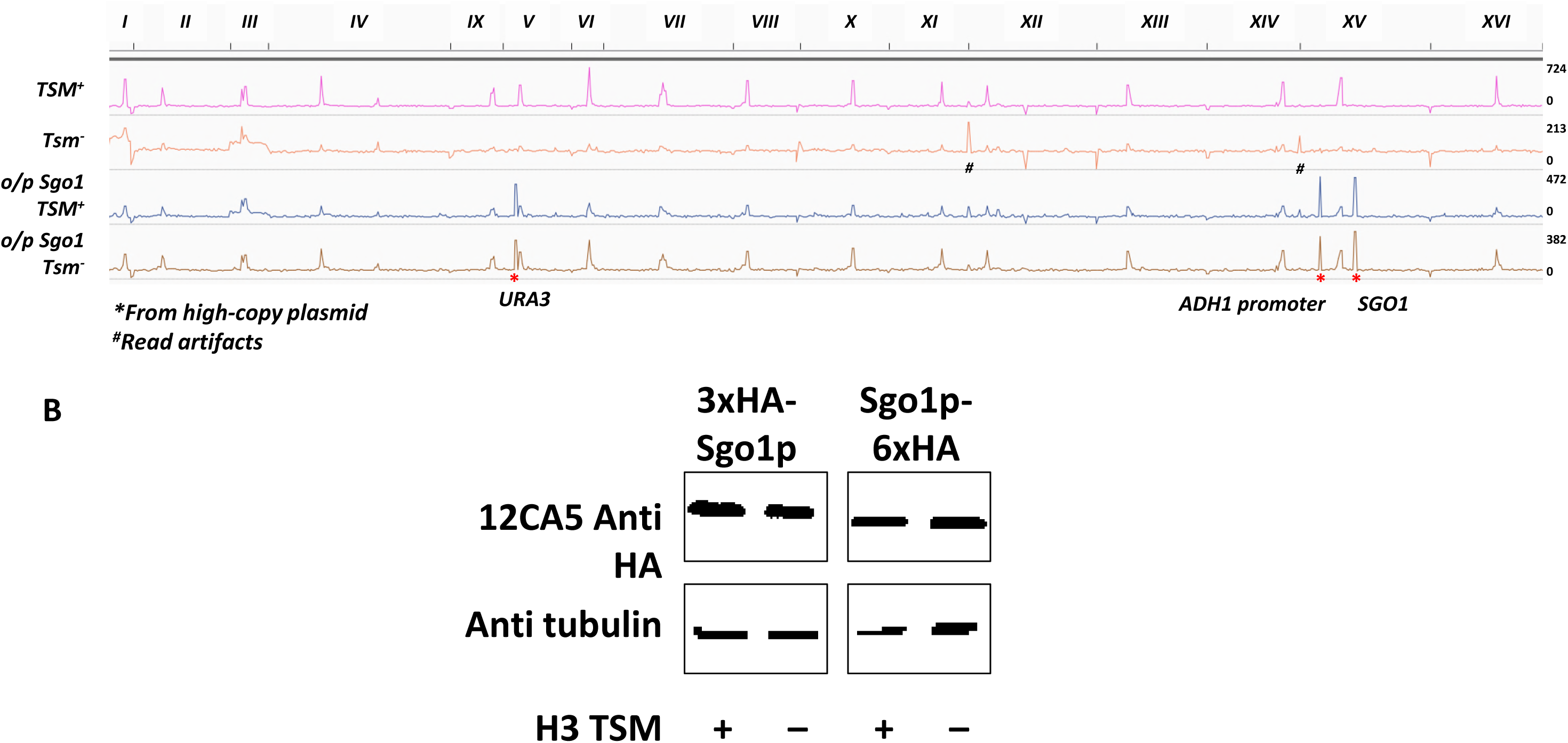
Sgo1p is localized only to the centromeric area in each chromosome. A. ChIP-seq data of Sgo1p expressed from its native locus or from a high-copy plasmid (o/p), in a wildtype or a tension sensing motif G44S mutant background, are presented as one single linear DNA. The range of each chromosome is shown on the top. B. Expression of Sgo1p is not affected by the status of the tension sensing motif. Sgo1p is tagged with HA at N’ or C’ terminus. The 3xHA-Sgo1p is expressed from a plasmid. The Sgo1p-6xHA is expressed from the native *SGO1* locus.

**Supplemental Figure 2.**
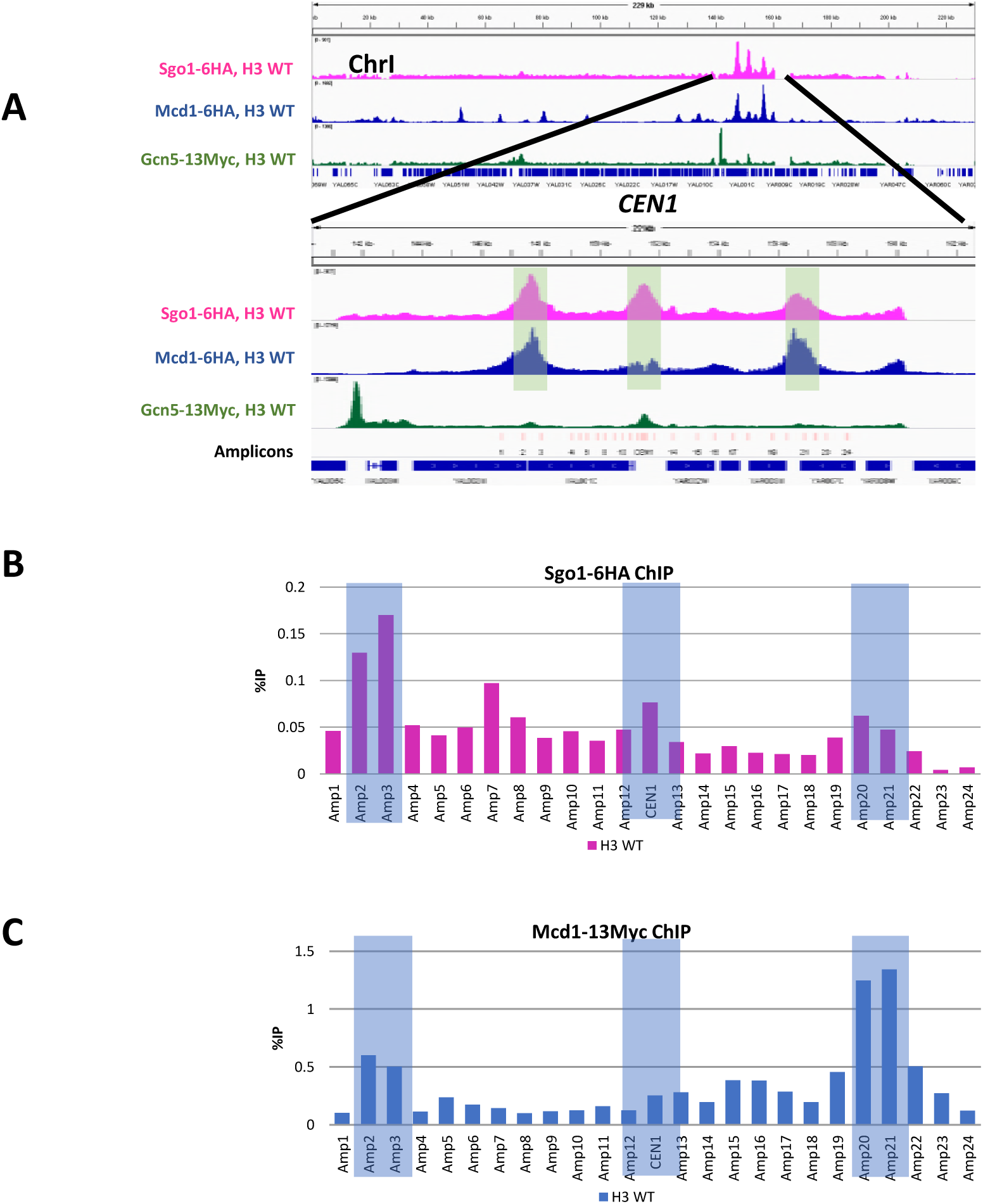
ChIP-qPCR to verify the ChIP-Seq findings. Shown are 25 amplicons spanning *CEN1*.

**Supplemental Figure 3.**
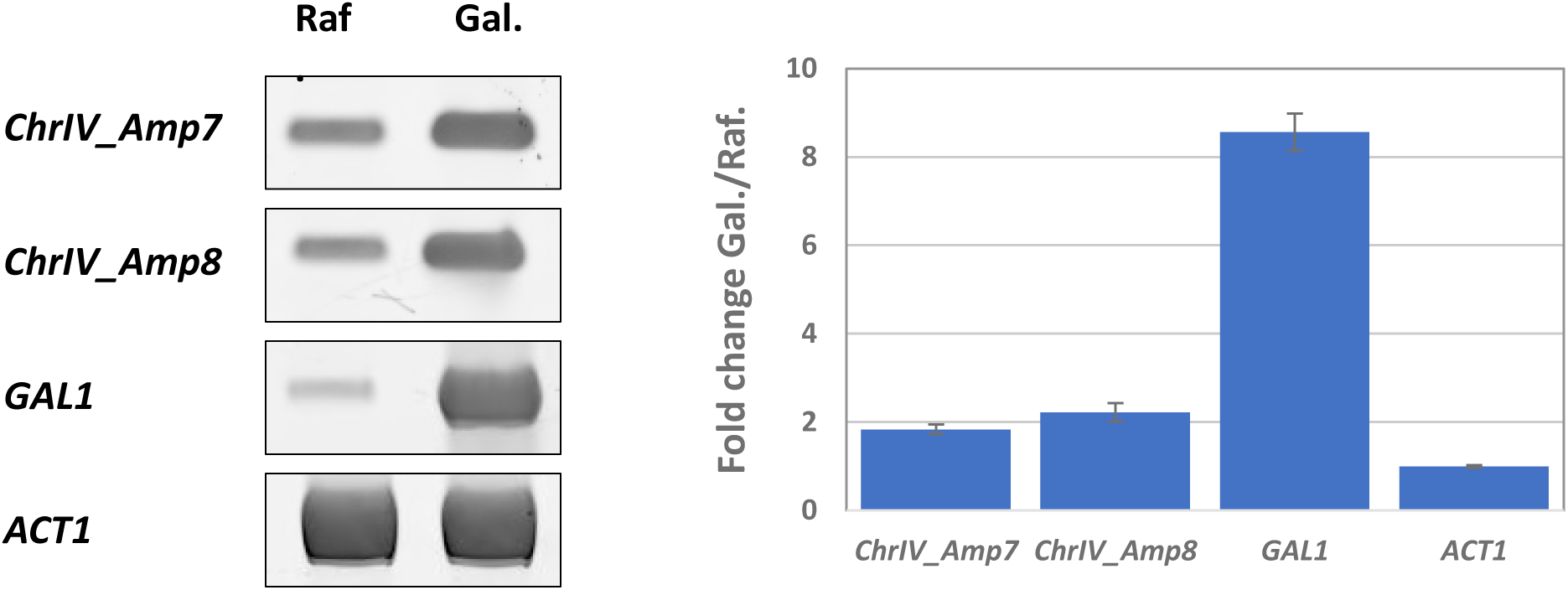
Ectopic, anti-sense expression of *YDR004W* from the *GAL1* promoter that replaces the cohesin associating region (CAR) at the 3’ end of *YDR004W*. Left panel shows the DNA gel images for reverse-transcription quantitative PCR of the indicated regions. ChrIV_Amp7 and 8 are within the *YDR004W* gene. *GAL1* and *ACT1* are positive and internal control for galactose induction. Right panel shows the quantification data, using ACT1 expression difference (Raf. vs. Gal.) for normalization. *YDR004W* is induced 2-fold by galactose.

**Supplemental Figure 4.**
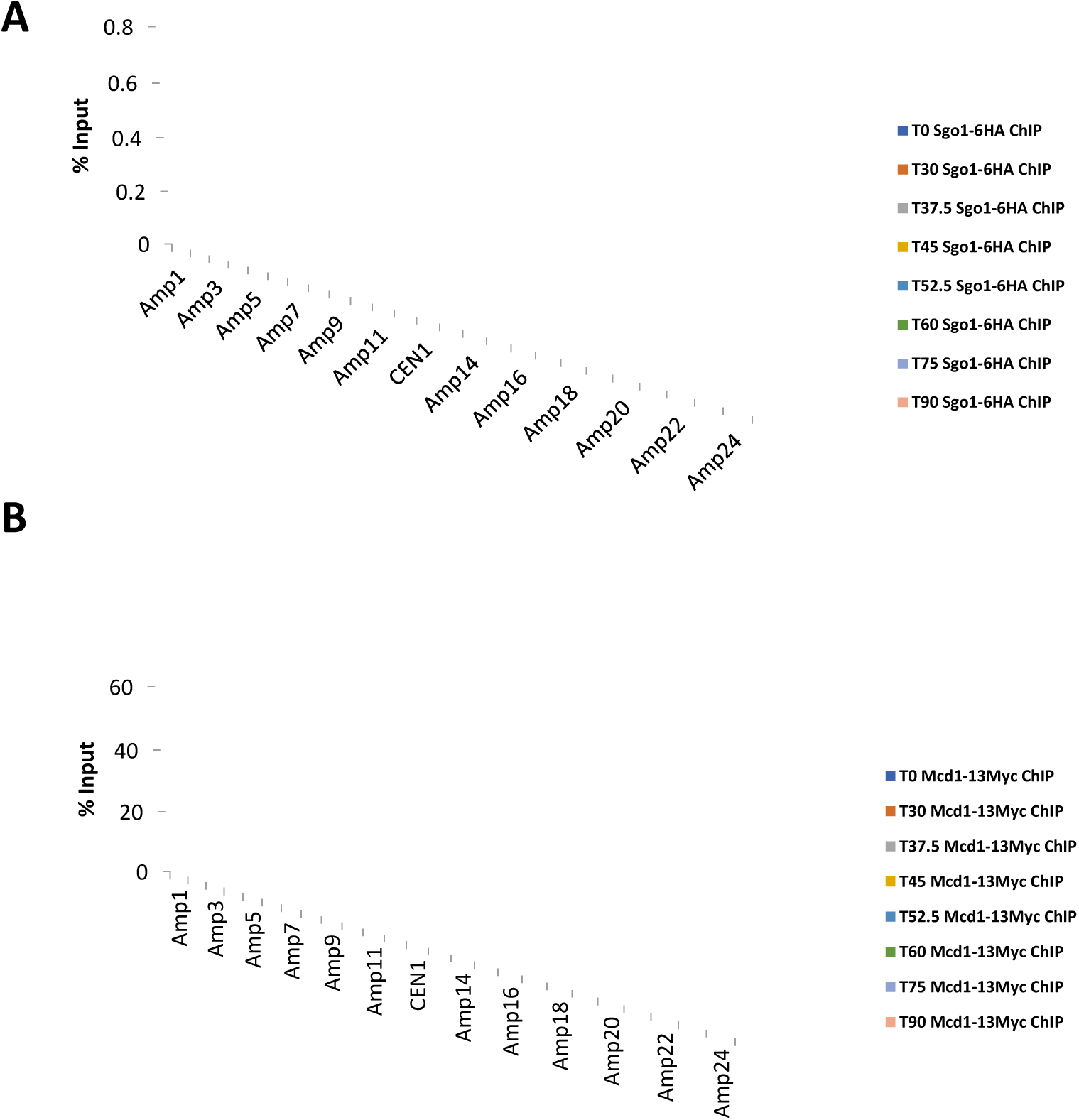
Dynamics of Sgo1p (A) and Mcd1p (B) localization at *CEN1* region. See **Figure 7** legends for description.

